# Senescence of alveolar stem cells drives progressive pulmonary fibrosis

**DOI:** 10.1101/820175

**Authors:** Changfu Yao, Xiangrong Guan, Gianni Carraro, Tanyalak Parimon, Xue Liu, Guanling Huang, Harmik J. Soukiasian, Gregory David, Stephen S. Weigt, John A. Belperio, Peter Chen, Dianhua Jiang, Paul W. Noble, Barry R. Stripp

## Abstract

Tissue fibrosis is a common pathological outcome of chronic disease that markedly impairs organ function leading to morbidity and mortality. In the lung, idiopathic pulmonary fibrosis (IPF) is an insidious and fatal interstitial lung disease associated with declining pulmonary function. Here, we show that alveolar type 2 (AT2) stem cells isolated from IPF lung tissue exhibit characteristic transcriptomic features of cellular senescence. We used conditional loss of Sin3a in adult mouse AT2 cells to initiate a program of p53-dependent cellular senescence, AT2 cell depletion, and spontaneous, progressive pulmonary fibrosis. We establish that senescence rather than loss of epithelial stem cells serves as a proximal driver of Tgfβ activation and progressive fibrosis and show that either genetic or pharmacologic interventions targeting p53 activation, senescence, or downstream Tgfβ activation, block fibrogenesis.

## Introduction

Fibrosis is a progressive form of pathological tissue remodeling seen following repeated injury or in the setting of chronic disease (1, 2). Although normal scarring is a physiologic and often reversable sequela of injury in most tissues, pathologic fibrosis of numerous organs, including lung, liver, and kidney, can be unrelenting and markedly impair function leading to organ failure. For example, patients with idiopathic pulmonary fibrosis (IPF) have a particularly poor prognosis with treatment options limited currently to medications that slow loss of lung function and when disease progresses, lung transplantation (3). Mechanisms involved in the initiation and progression of fibrosis, such as that seen in patients with IPF, are poorly defined and most likely multifactorial. However, common features of IPF, which mirror disease manifestation in many organs, are unresolved healing of wounded epithelium, activation and proliferation of (myo)fibroblasts and invasive fibroblasts (4-6), excessive deposition of extracellular matrix, particularly fibrillar collagens, and destruction of lung architecture leading to declining pulmonary function and death (3, 7-9). Recent studies have implicated mechanisms of accelerated aging, including loss of epithelial progenitor cell function and/or numbers and cellular senescence, in IPF pathogenesis (10-13). Indeed, in about 20% of familial IPF cases, mutations have been found in genes involved in telomere function (TERT and TERC) and protein folding and secretion that impact the function of epithelial cells (14-16). Furthermore, disease-associated gene variants defined in genome-wide screens have been linked to defects in host defense and regulation of cellular senescence (2, 17).

Normal maintenance of the alveolar epithelium is accomplished through the proliferation of alveolar type 2 (AT2) cells, a facultative stem cell that produces and secretes pulmonary surfactant in its quiescent state (18). However, stress from genetic or environmental factors can compromise the ability of AT2 cells to transition from quiescent to proliferative states, thereby leading to defective epithelial maintenance (19). We recently demonstrated that fibrotic regions of lung tissue from IPF patients show both regional depletion of AT2 cells and the presence of epithelial cells with abnormal activation of multiple pathways, including p53 and TGFβ signaling (20). In other organs, such changes in cell phenotype promote cellular senescence (21-31). Thus, age-dependent senescence of AT2 cells, accompanied by loss of stem/progenitor cell function, could contribute to progressive lung fibrosis.

Here, we define the senescence-associated phenotype of AT2 cells in lung tissue of patients with IPF and establish a novel mouse model to test further the contribution of AT2 cell dysfunction in progressive lung fibrosis. To this end, we generated mice with AT2-specific conditional loss of Sin3a, a key component of Sin3-HDAC complex that regulates chromatin structure and gene expression in all eukaryotic cells. Silencing of Sin3a led to defective progenitor cell function and induction of a program of p53-dependent AT2 senescence. In addition, we show that senescence of AT2 cells is sufficient to initiate TGFβ-dependent progressive lung fibrosis that closely resembles pathological remodeling seen in IPF lungs. Fibrosis was diminished either by selective loss of p53 function in AT2 cells, through systemic inhibition of TGFβ signaling, or ablation of senescent cells by systemic delivery of senolytic drugs. Our data suggest that p53-induced AT2 senescence serves as a proximal driver and candidate therapeutic target in progressive lung fibrosis and provides key evidence to support roles for epithelial dysfunction in both initiation and progression of distal lung fibrosis seen in IPF.

## Results

### Senescent phenotype of AT2 cells in end-stage IPF lung tissue

To define senescent cell types of the IPF lung, we evaluated the expression of genes linked to the senescence-associated secretory phenotype (SASP) (32, 33) by mining the Lung Genomics Research Consortium (LGRC) transcriptomic database (http://www.lunggenomics.org). Our analyses revealed a large proportion of known senescence associated genes including SASP genes whose transcripts were significantly elevated in ILD lungs compared to normal control tissue (Fig 1A). Furthermore, staining of IPF explant tissue for senescence-associated β-gal (SA-βgal) revealed multiple senescent cell types including epithelial cells (Fig 1B). To test the contribution of alveolar Type II (AT2) cells to the senescent cell fraction, we took advantage of RNA-Seq data generated from enriched HTII-280+ epithelial cells isolated from either control donor lung or IPF explant lung tissue (20). As with the LGRC data set, bulk RNA-seq data showed increased expression of senescence associated genes including SASP genes in HTII-280+ AT2 cells isolated from IPF patient samples compared to control donors (Fig 1C). These data demonstrate that AT2 cells contribute to the total pool of senescent lung cells in IPF.

**Figure 1.**
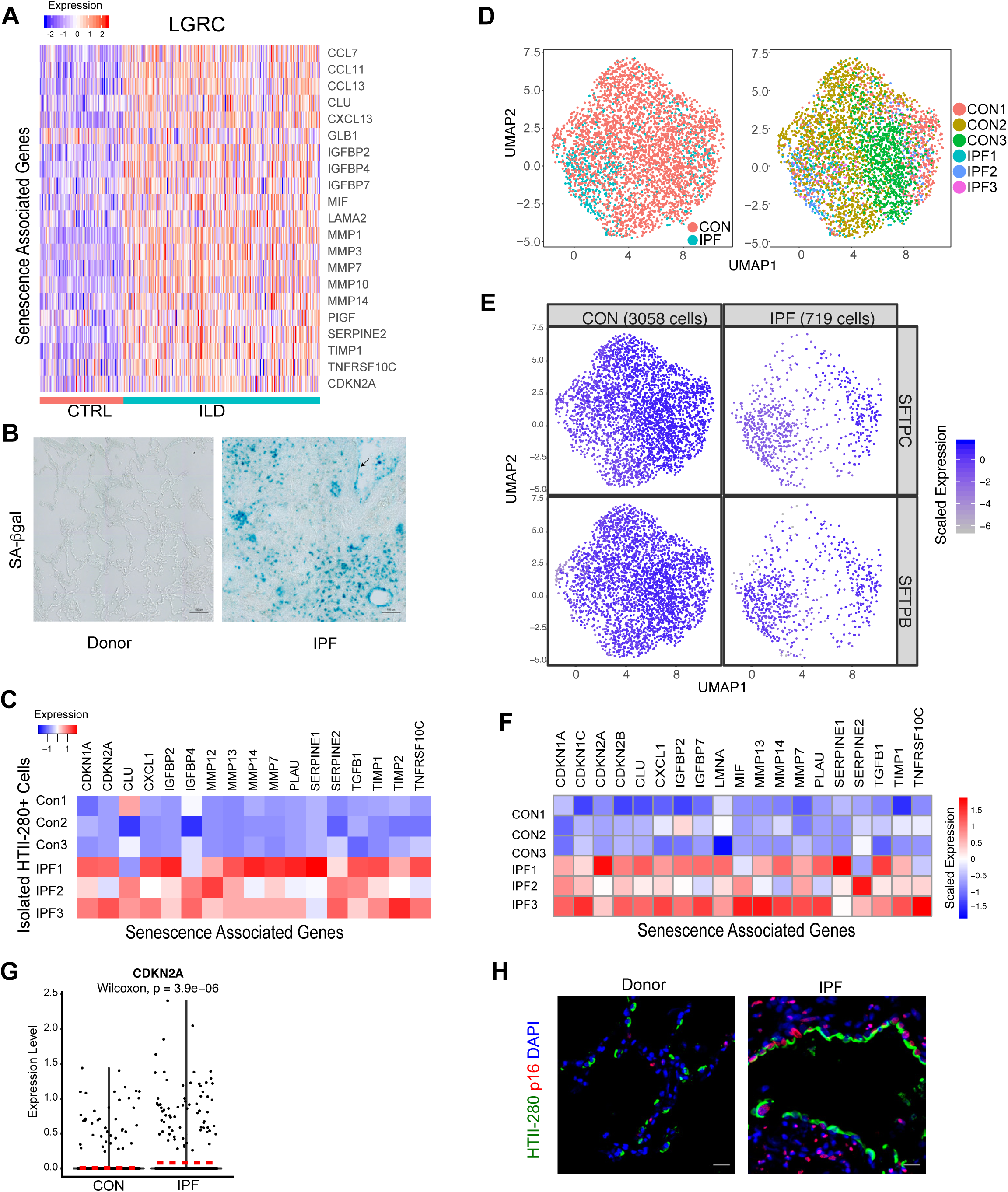
Accumulation of senescent AT2 cells in IPF explant tissue. A. Heatmap of known SASP genes derived from the LGRC data set comparing ILD patient samples (253) with control (CTRL) samples (108). B. SA-βgal signaling of donor and IPF lung sample. Arrow indicates possible epithelial cell layer. C. Heatmap of known SASP genes derived from bulk RNA-Seq of isolated human HTII-280 cells from control donor and IPF patient samples. D. UMAP visualization of AT2 cell subsets, origin and clustering, from human epithelial cell scRNA-Seq. E. UMAP visualization showing relative expression of SFTPC in AT2 cells comparing control and IPF, from human epithelial cell scRNA-Seq. F. Heatmap of known SASP genes and senescence biomarkers for average expression of these genes from each patient sample (AT2 cell subsets from human epithelial cell scRNA-Seq). G. Violin plot representation showing relative expression of CDKN2A/p16 in human scRNA-Seq AT2 cells subset. Dash line indicates median expression of each group. H. HTII-280 and CDKN2A/p16 immunostaining on control donor lung distal tissue and IPF fibrotic tissue. Scale bar = 20μm.

To further validate these observations, we performed single cell RNA-sequencing (scRNA-Seq) with fluorescence-activated cell sorting (FACS) enriched epithelial cells (DAPI^-^, CD45^-^, CD31^-^, CD326^+^) from either control donor distal lung tissues or IPF fibrotic lung tissues (Feature H&E Supplemental Figure 1A, 1B). A total of 10146 captured single cells passed quality control for further batch correction and unsupervised clustering (Supplemental Figure 2A). Major cell types, including AT1, AT2, club, ciliated, goblet, and basal cells, were assigned according to their expression of known cell type-specific gene signatures and visualized by Uniform Manifold Approximation and Projection (UMAP) (Supplemental Figure 2A, B). AT2 cells were identified and selected for further analysis based upon their clustering and abundant expression of transcripts for SFTPC and SFTPB (Fig 1D, E). Expression of both SASP-related genes and other senescence biomarkers, including CDKN1A/p21, CDKN2A/p16 were significantly elevated in single cell transcriptomes of IPF AT2 cells compared to control AT2 cells (Fig 1F, 1G and also 4F). These findings were recapitulated among single AT2 cell transcriptomes evaluated in a distinct data set of control and IPF subjects (34), available in gene expression omnibus (GSE122960; Supplement Fig 2C-E). Further validation by immunofluorescence staining revealed increased CDKN2A/p16 and CDKN1A/p21 immunoreactivity in HTII-280+ AT2 cells within localized hyperplastic epithelium adjacent to fibrotic regions of IPF lung tissue (Fig 1G, also Fig 4E). We also assessed the contribution of AT2 cell senescence in the development of lung fibrosis using intratracheal administration of bleomycin, a commonly used animal model of lung injury and fibrosis (35). We evaluated our scRNA-Seq analysis comparing cells from mouse lungs at day-21 after bleomycin treatment – when fibrosis peaks – with vehicle-treated litter mate controls(36). We analyzed AT2 cells for SASP factor genes expression and found increased transcript levels of SASP genes in AT2 cells from bleomycin-treated compared to cells from control mice (Supplemental 3 Fig A-E). Together, these data demonstrate that senescence of AT2 stem cells is a common pathological feature of IPF observed among all patient samples regardless of genetic and other initiating factors.

**Figure 2.**
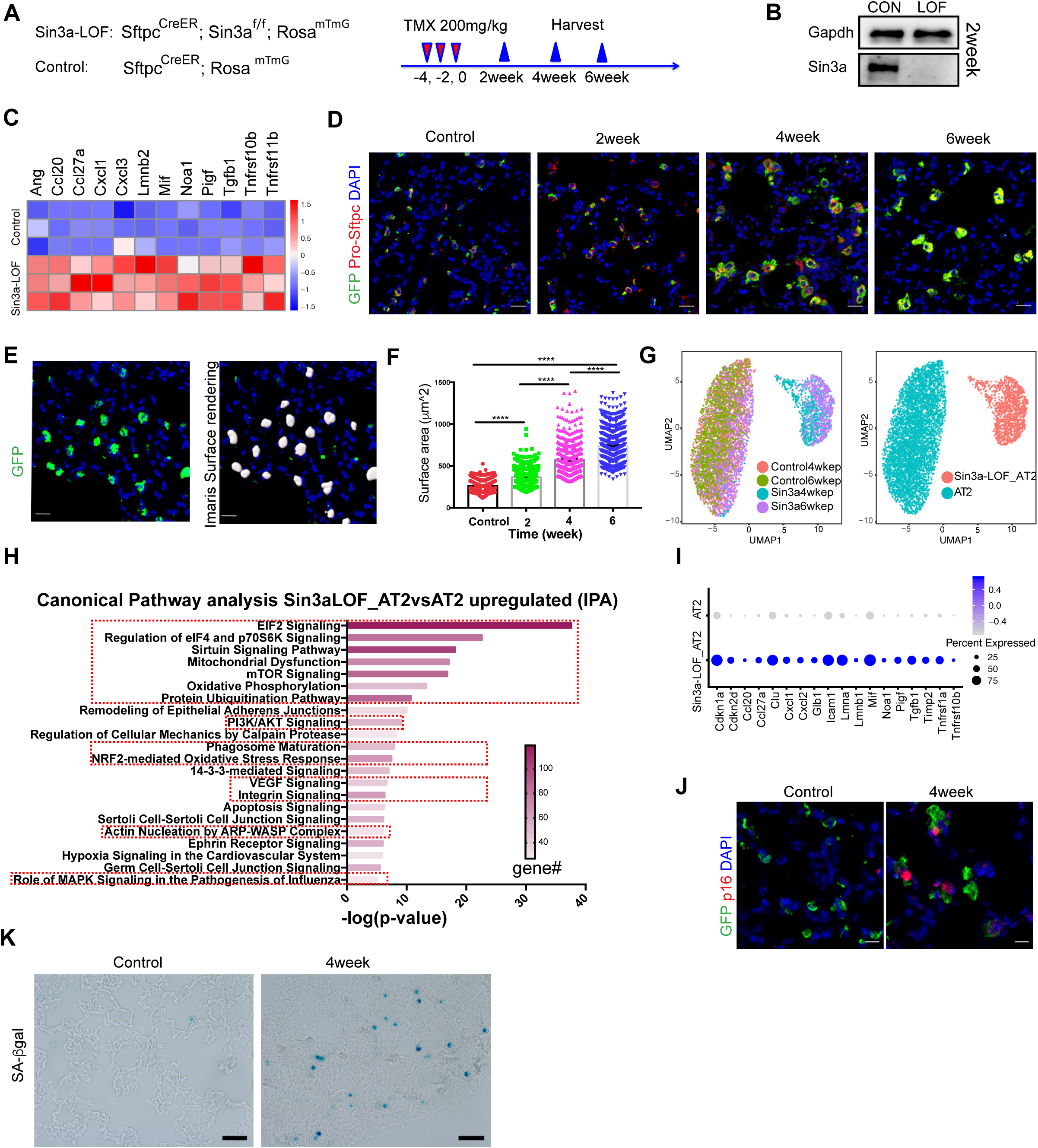
Loss of Sin3a in adult mouse AT2 cell leads to cellular senescence. A. Schematic outline of experiment design. B. Western Blot detection of Sin3a knockout efficiency in isolated AT2 cells 2 weeks after tamoxifen treatment. C. Heatmap of SASP genes from RNA-Seq of isolated AT2 cells 2 weeks after tamoxifen treatment. D. Time course of immunofluorescence staining for lineage reporter (GFP) and Pro-Sftpc in control and Sin3a-LOF lung. Scale bar = 20μm. E. Representative immunofluorescence staining of membrane-bound GFP and Imaris surface rendering for surface area quantification. F. AT2 cell surface area quantification. G. UMAP visualization of AT2 cell subset, origin and clustering, from mouse epithelial cell scRNA-Seq. H. IPA canonical pathway analysis of genes up-regulated in AT2 cells of Sin3a-LOF compared to control; senescence related pathways are highlighted. I. Expression of SASP genes and senescence biomarkers between Sin3a-LOF AT2 cells and control AT2 cells. J. Immunofluorescence staining of GFP and Cdkn2a/p16 of control and Sin3a-LOF mouse lung. Scale bar = 10μm. K. SA-βgal signaling of control and Sin3a-LOF mouse lung (4 weeks post-tamoxifen). p-value calculated by two-tailed student t-test. * p<0.05, ** p<0.01, *** p<0.001, **** p<0.0001.

**Figure 3.**
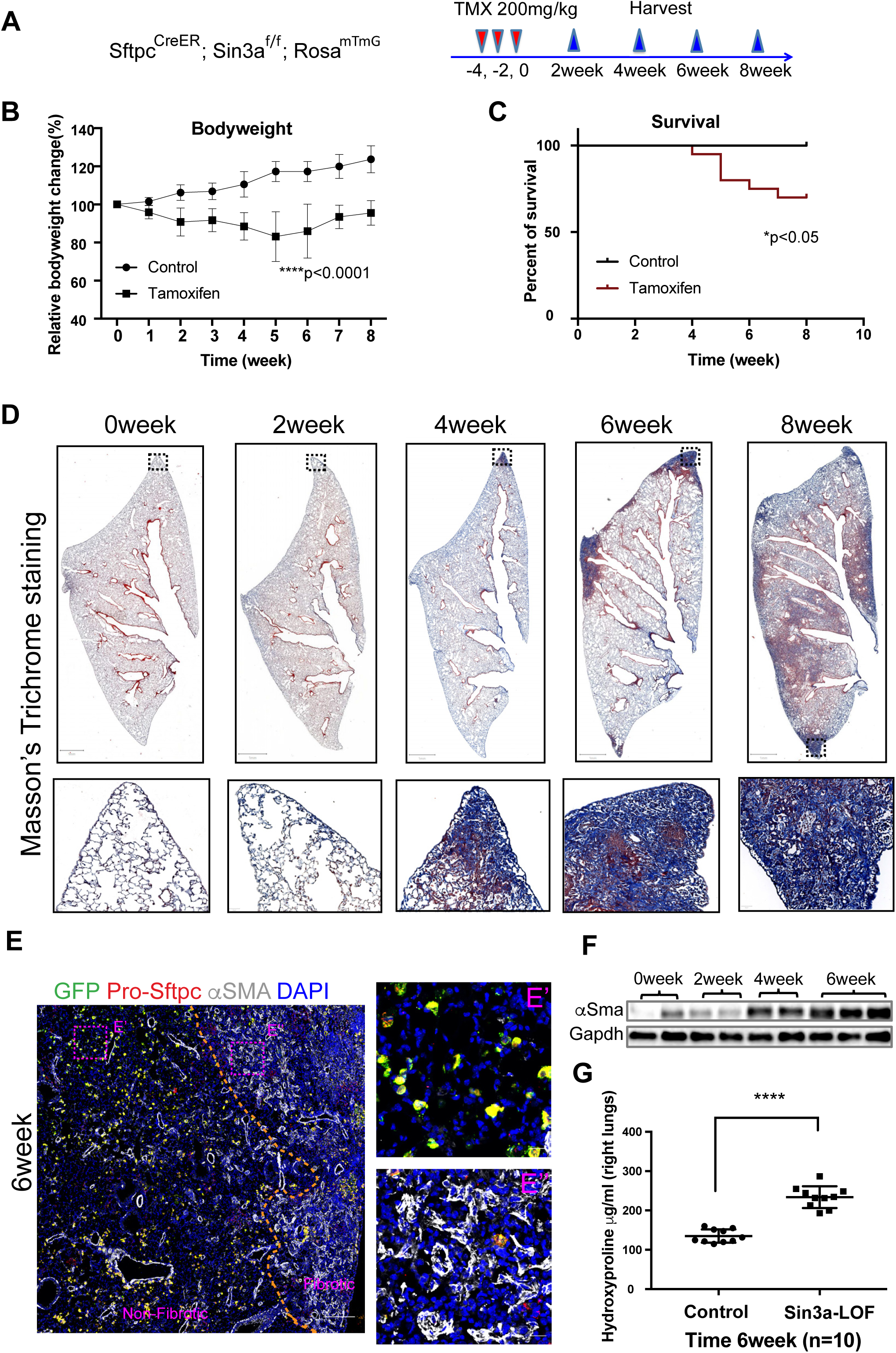
Loss of Sin3a in AT2 cells leads to progressive lung fibrosis. A. Schematic outline of experiment design. B. Bodyweight change for tamoxifen-exposed Sin3a-LOF and control mice. p<0.0001 for two-way ANOVA test and two-tailed student t-test comparing Control vs Tamoxifen treated group at each time point 2 weeks after tamoxifen exposure. C. Survival curve for tamoxifen-exposed Sin3a-LOF and control mice. *p<0.05 by Log-rank (Mantel-Cox) survival analysis. D. Time course of Masson’s trichrome staining of lung tissue from tamoxifen-exposed Sin3a-LOF and control mice. E. Representative immunofluorescence staining of Sin3a-LOF lung tissue 6 weeks post-tamoxifen exposure for lineage reporter gene (GFP), Pro-Sftpc and αSMA. Dash line represents the boundary between fibrotic and non-fibrotic tissue. E’ and E’’, zoom images of representative regions from either non-fibrotic or fibrotic tissue. Scale bar = 200μm and 20μm, respectively. F. Western Blot detection of αSMA abundance within lung tissue of Sin3a-LOF mice. G. Hydroxyproline content of right lung between Sin3a-LOF and control mice 6 weeks post-tamoxifen exposure (n=10 for each group). p-value calculated by two-tailed student t-test. * p<0.05, ** p<0.01, *** p<0.001, **** p<0.0001.

**Figure 4.**
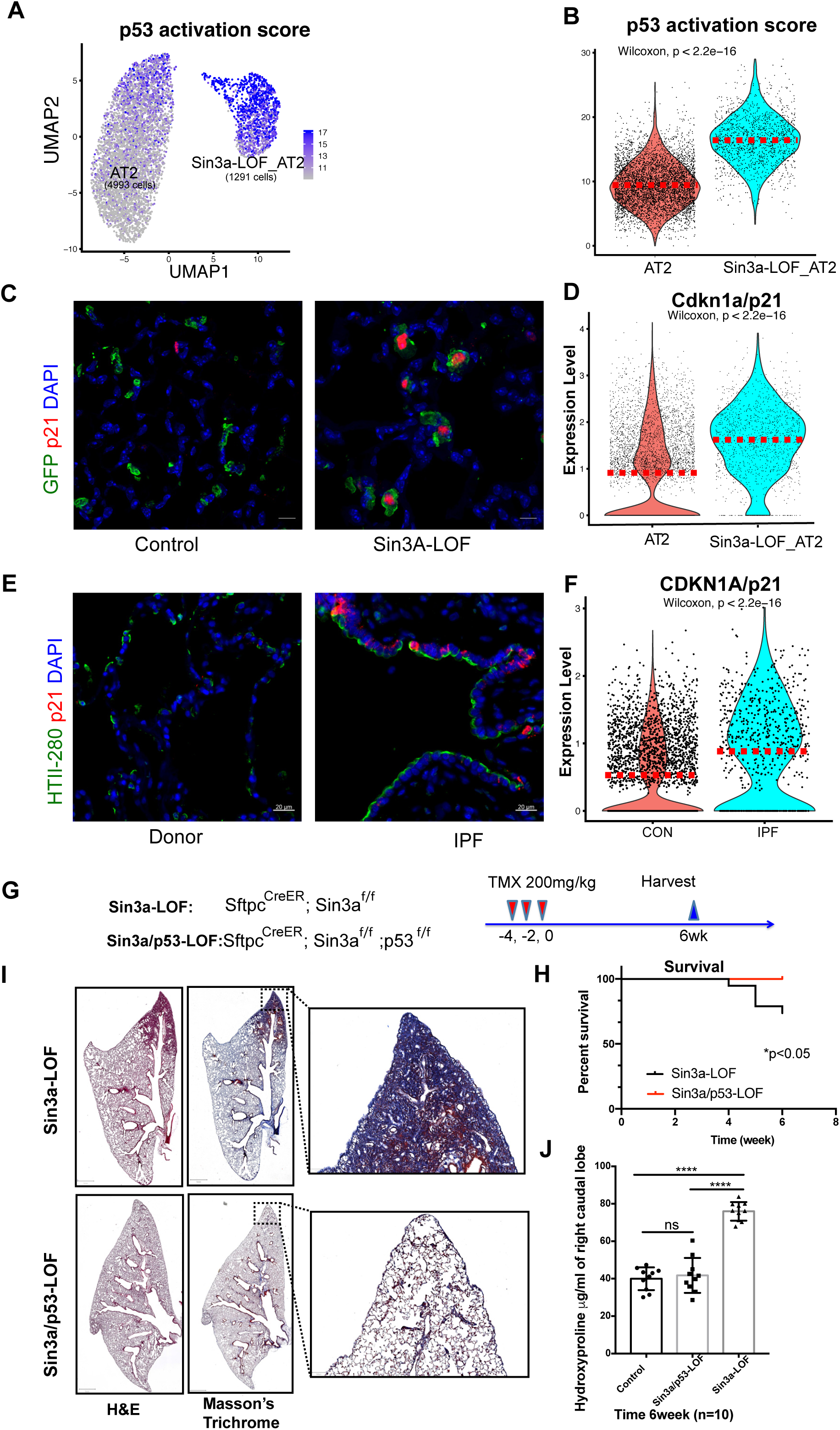
p53 pathway activation in Sin3a-LOF AT2 cells. A. UMAP visualization of p53 activation score calculated using p53 KEGG pathway genes of mouse scRNA-Seq AT2 cells subset. B. Violin Plot visualization of p53 activation score of mouse scRNA-Seq AT2 cells subset. Dash line indicated median expression of each group. C. Representative immunofluorescence staining of lineage reporter gene (GFP) and Cdkn1a/p21, comparing control lung tissue with Sin3a-LOF 6 weeks post-tamoxifen exposure. Scale bar = 10μm. D. Violin plot representation showing relative expression of Cdkn1a/p21 in mouse scRNA-Seq AT2 subset. Dash line indicates median expression of each group. E. Representative immunofluorescence staining of HTII-280 and CDKN1A/p21, comparing control donor distal lung tissue with IPF patient fibrotic lung tissue. Scale bar = 20μm. F. Violin plot representation showing relative expression of CDKN1A/p21 in human scRNA-Seq AT2 cells subset. Dash line indicates median expression of each group. G. Schematic outline of experiment design for p53-LOF studies. H. Survival curve for Sin3a-LOF and Sin3a/p53-LOF groups 6 weeks post-tamoxifen treatment. *p<0.05 by Log-rank (Mantel-Cox) survival analysis. I. Hematoxylin & Eosin (H&E) and Masson’s trichrome staining of Sin3a-LOF and Sin3a/p53-LOF lung tissues 6 weeks post-tamoxifen treatment. J. Hydroxyproline content of right caudal lobe in Sin3a-LOF and Sin3a/p53-LOF mice 6 weeks post-tamoxifen treatment (n=10 for each group). p-value calculated by two-tailed student t-test. * p<0.05, ** p<0.01, *** p<0.001, **** p<0.0001.

### A novel mouse model of conditional AT2 stem cell senescence

We next sought to determine whether senescence of AT2 cells drives progressive fibrosis. In our previous work, we demonstrated that endodermal progenitor cells of the developing mouse lung are uniquely dependent upon Sin3a and that loss of Sin3a in developmental endodermal progenitor cells leads to senescence and aberrant lung development (37). To determine if the progenitor function of adult lung AT2 cells have a similar dependence on Sin3a, we generated Sftpc^CreER^; Sin3a^f/f^; Rosa^mTmG^ to allow tamoxifen-induced conditional Sin3a loss-of-function (LOF) in AT2 cells (herein referred to as Sin3a-LOF; Fig 2A). Efficiency of Sin3a loss was verified by immunoblotting of homogenates of isolated AT2 cells (Fig 2B) and by immunostaining of lung tissue sections (Supplemental Figure 4A). Both methods demonstrated near complete loss of Sin3a within AT2 cells 2 weeks after tamoxifen administration. Furthermore, Sin3a-LOF led to about an 80% reduction in AT2 colony forming efficiency when isolated AT2 cells were placed in 3D culture (Supplemental Figure 5E and F) indicating impaired progenitor capability of AT2 cells due to loss of Sin3a.

To better define how loss of Sin3a impacts the phenotype of AT2 cells, we performed bulk RNA-Seq on AT2 cells isolated from lungs of Sin3a-sufficient (wildtype) and Sin3a-LOF mice. Using DEseq2 test with an adjusted p-value cut off of 0.05, we found that 3489 transcripts were significantly upregulated and 3605 transcripts were significantly downregulated in Sin3a-LOF AT2 cells compared to wildtype cells (Supplemental Figure 4B, C). We then used Ingenuity Pathway Analysis (IPA) to identify differentially regulated gene networks within Sin3a-LOF AT2 cells compared to their Sin3a-sufficient counterparts. Upregulated pathways in Sin3a-LOF AT2 cells included oxidative phosphorylation, mitochondrial dysfunction, sirtuin signaling, EIF2 signaling, and mTOR signaling. Interestingly, these pathways are all linked to cellular senescence(38-53). Conversely, the most highly downregulated pathway was linked to maintenance of embryonic stem cell pluripotency (Supplemental Figure 4D, E). Furthermore, we also identified a number of SASP genes, including Ang, Cxcl1, Cxcl3, and Mif, whose transcript levels were upregulated in Sin3a-LOF AT2 cells (Fig 2C). These changes in the molecular phenotype of AT2 cells were accompanied by dramatic increases in the size of AT2 cells (Fig 2D-F and Supplemental Figure 4A), a characteristic feature of senescent cells (54, 55), Indeed, at 6 weeks post-tamoxifen treatment, the average cross-sectional surface area of AT2 cells increased almost 3-fold in Sin3a-LOF mice (735.5 ± 8.076 μm^2^) compared to age-matched Sin3a-sufficient controls (263 ± 2.797 μm^2^) (Fig 2F).

To define further the molecular changes accompanying Sin3a loss within AT2 cells, we performed single cell RNA sequencing (scRNA-Seq) on enriched epithelial cells (DAPI^-^, CD45^-^, CD31^-^, CD326^+^) isolated from Sin3a-LOF and control mouse lungs 4 weeks and 6 weeks after tamoxifen treatment. A total of 11686 captured single cells passed quality control for further batch effect correction and unbiased clustering. Molecular similarity between cells was visualized by t-Distributed Stochastic Neighbor Embedding (t-SNE), with major cell types assigned to clusters according to expression of known cell type-specific gene signatures and the Cre-activated GFP lineage reporter (Supplemental Figure 5A, B). AT2 cells were defined by expression of Sftpc and GFP transcripts were selected for further analysis and formed two distinct clusters when visualized by UMAP (Fig 2G). The largest of these AT2 clusters included representation from both Sin3a-sufficient and Sin3a-LOF mice recovered at 4 or 6 weeks after tamoxifen treatment. In contrast, the smaller AT2 cluster only included cells derived from Sin3a-LOF mice and were considered to represent the bulk of AT2 cells having recombined both Sin3a-flox alleles to generate Sin3a-LOF AT2 cells. Using Wilcoxon rank sum test with a cut off of p-value = 0.01, we identified 2683 genes that were significantly upregulated and 518 genes that were significantly downregulated in Sin3a-LOF AT2 cells compared to Sin3a-sufficient AT2 cells (Supplemental Figure 5C).

We then used IPA to perform canonical pathway analysis on our scRNA-Seq data to assess the potential for these changes in gene expression to influence cellular phenotype. Pathways up-regulated pathways in Sin3a-LOF AT2 cells compared to wildtype AT2 cells included oxidative phosphorylation, mitochondrial dysfunction, sirtuin signaling, EIF2 signaling, and mTOR signaling, which were all identified in our bulk AT2 cell RNA-seq analyses. In addition, our scRNA-Seq data revealed differential expression of several other genes and pathways linked to cellular senescence, including eIF4 and p70S6K (46), protein ubiquitination (56, 57), PI3K/AKT signaling (21-24), phagosome maturation (58, 59), VEGF signaling (60), integrin signaling (61), NRF2-mediated oxidative stress (62-65), actin nucleation (66), and MAPK signaling (67-69) (Fig 2H). Furthermore, increased expression of SASP genes, as well as senescence biomarkers including Cdkn1a/p21, Trp53, and others, were prominent in Sin3a-LOF AT2 cells compared to Sin3a-sufficient AT2 cells (Fig 2I). We used immunofluorescence to demonstrate expression of senescence-associated cell cycle inhibitors Cdkn2a/p16 and Cdkn1a/p21 co-localized with the GFP lineage trace (AT2 cells). Cdkn2a/p16 and Cdkn1a/p21 co-localized with GFP used to traced AT2 cells. Immunofluorescence signal for both cell cycle regulators was dramatically increased within GFP+ AT2 cells of Sin3a-LOF mice compared to their signal in wildtype cells (Fig 2J and Fig 4C). Increased expression of p16 and p21 was also accompanied by an increase in senescence-associated β–gal staining in Sin3a-LOF mouse lung samples compared to wildtype lung (Fig 2K). Together, these data demonstrate that loss of Sin3a in adult AT2 cells activates cellular senescence.

### Sin3a-LOF and associated senescence of AT2 cells leads to progressive lung fibrosis

Having found that Sin3a-LOF induces senescence within AT2 cells, we next sought to determine if AT2 cell dysfunction mediated by Sin3a-LOF impacted lung fibrosis. 10-12-week-old Sin3a-LOF mice were exposed to three doses of tamoxifen. Lung injury and remodeling were assessed over the following 8 weeks (Fig 3A). Sin3a-LOF led to persistent morbidity, as indicated by a decline in bodyweight, and a significant increase in mortality (either spontaneous death or reaching euthanasia criteria), both of which were not seen in control mice (Fig 3B and C). Histological assessment revealed focal consolidations of matrix deposition that initiated at 4 weeks at the lung edges and progressed inward forming extensive interstitial fibrosis at later times (Fig 3D). Importantly, this pattern of fibrosis mirrors closely both the onset and progression of fibrosis seen in human lung (70, 71).

We performed RNA-Seq of whole lung RNA to define global changes in gene expression that accompany progressive fibrosis in Sin3a-LOF mice. Complementing our histologic findings, IPA identified altered pathways controlling cell proliferation and activation of fibrosis at 4 weeks post-tamoxifen, which increased in significance at later times (Supplemental Figure 6A). Paralleling the increases in fibrosis gene expression scores was a marked stimulation of multiple collagen and other fibrosis-related genes, such as fibronectin, Ctgf (Supplemental Figure 6B, C), SASP-related genes (Supplemental Figure 6D), and the myofibroblast marker αSMA (Fig 3 E, F). An elevated signal for αSMA immunoreactivity was seen in fibrotic regions (Fig 3E). In addition, hydroxyproline content, a marker of collagen deposition, was markedly elevated in Sin3a-LOF lungs compared to control tissue (Fig 3G). Collectively, these data demonstrate that AT2 cell dysfunction and senescence resulting from loss of Sin3a is sufficient to drive progressive lung fibrosis in a pattern that mimics closely the pattern of fibrosis seen in IPF patients.

### p53 signaling pathway activation in senescent AT2 cells

We reported that p53 and TGFβ signaling are dramatically upregulated within AT2 cells isolated from explant IPF tissue (20). We used IPA analysis of total lung RNA-seq data to identify candidate regulators of fibrosis in Sin3a-LOF mice. TGFB1 (highlighted in red) and TP53 (highlighted in blue) were identified as the most differentially expressed upstream regulators (Supplemental Figure 7A). Similarly, we also identified TGFB1 (highlighted in red) and TP53 (highlighted in blue) by IPA upstream analysis using 1) bulk RNA-seq of isolated HTII-280+ cells from IPF vs control lungs; 2) scRNA-Seq of human lungs comparing IPF vs control AT2 cells; and 3) scRNA-Seq of murine lungs comparing Sin3a-LOF vs control AT2 cells (Supplemental Figure 7B-7D). Thus, these data indicate that TGFB1 and TP53 are critical regulators of pro-fibrotic pathway in both human and mouse epithelial samples.

Given that p53 activation is linked to a loss of progenitor cell function and induction of cellular senescence (72), we evaluated p53 activation status in the Sin3a-LOF mouse model and human IPF patient samples. As discussed above, we found that p53 signaling was among the top upregulated pathways in lung tissue of Sin3a-LOF mice (Supplemental Figure 5A). These findings were confirmed by immunoblotting, which showed that p53 protein levels increased markedly and progressively following tamoxifen exposure in Sin3a-LOF mice (Supplemental Figure 7E). Increases in p53 protein abundance were also evident 7 and 14 days after exposure of mice to intratracheal bleomycin (Supplemental Figure 7F).

To assess p53 pathway activation within individual lung AT2 cells, we compared the p53 pathway activation gene list from the KEGG database to our scRNA-Seq data generated with AT2 cells from Sin3a-LOF mice and with human AT2 cells from IPF tissue. We observed a significant increase in p53 activation score both in Sin3a-LOF AT2 cells versus Sin3a-sufficient control AT2 cells (Fig 4A, B) and in IPF AT2 cells versus control human AT2 cells (Supplemental Figure 7G). These findings were further verified by gene expression analysis of Cdkn1a/p21, a known downstream target of activated p53 (72, 73). Compared to Sin3a-sufficient controls, increased immunoreactivity for Cdkn1a/p21 was observed in GFP+ AT2 cells in lung tissue of Sin3a-LOF mice (Fig 4C) and was accompanied by a significant increase in the proportion of AT2 cells with detectable Cdkn1a/p21 mRNA and in the median expression levels per cell (Fig 4D). Similarly, increased CDKN1A/p21 immunoreactivity and mRNA expression were observed in HTII-280+ AT2 cells from IPF lung tissue compared to these cells from control donor lung (Fig 4E and F, Supplemental Figure 7H).

To examine further the importance of p53 activation as a downstream mediator of progressive lung fibrosis in Sin3a-LOF mice, we generated Sftpc^CreER^; Sin3a^f/f^; p53^f/f^ mice (herein referred as Sin3a/p53-LOF) with conditional loss of both Sin3a and p53 in AT2 cells. 10-12-week-old Sin3a-LOF and Sin3a/p53-LOF mice were exposed to tamoxifen, and viability and lung injury monitored for 6 weeks (Fig 4G). In contrast to Sin3a-LOF mice, Sin3a/p53-LOF mice were protected from bodyweight decline and mortality, also accompanied by a decrease in senescence-associated β–gal staining in Sin3a/p53-LOF mouse lung samples compared to Sin3a-LOF mouse lung samples (Fig 4H and Supplemental Figure 7I, 7J). Consistent with a protective effect of p53-LOF, fibrosis seen in Sin3a-LOF mice was diminished in Sin3a/p53-LOF mice (Fig 4I-4J). We found similar results in Sin3a-LOF mice that received intraperitoneal injections of p53 inhibitor Pifithrin-α every other day after the first week of tamoxifen treatment (Supplemental Figure 8 A-E). These data demonstrate that silencing p53 expression or activity in AT2 cells prevented the pro-fibrotic phenotype that results from Sin3a-LOF and suggests that p53 activation represents a critical determinant of AT2 progenitor cell dysfunction leading to progressive lung fibrosis.

### Inhibition of TGFβ alleviates lung fibrosis due to Sin3a loss of function in AT2 cells

Activation of both p53 and TGFβ signaling was seen in AT2 cells of Sin3a-LOF mice and in lung tissue from patients with IPF (Supplemental Figure 7). TGFβ activation plays a key role in lung fibrosis (74-77), especially in epithelial cells (78, 79). Tgfb1 may be a key mediator of SASP (33). We next interrogated our scRNA-Seq data of Sin3a-LOF mouse model and human IPF specimens to determine if there was evidence of TGFβ induction in AT2 cells. The abundance of Tgfb1 and TGFB1 mRNAs were significantly increased both in Sin3a-LOF and IPF AT2 cells compared to their matched control AT2 cells (Fig 5A-C; Supplemental Figure 9A-C). TGFβ activation was also accompanied by a general decrease in abundance of SFTPC mRNA in IPF and Sin3a-LOF AT2 cells (Fig 1E, Supplemental Figure 2D, 2F, 2G; Fig 5E, F). Hence, Sin3a-LOF AT2 cells were subdivided into Sftpc^+^ and Sftpc^-^ subsets and compared to Sin3a-sufficient AT2 cells (Fig 5G). We found that Sin3a-LOF/Sftpc^-^ AT2 cells showed the highest TGFβ activation score, with Sin3a-LOF/Sftpc^+^ AT2 cells showing an intermediate score, and Sin3a-sufficient AT2 cells with the lowest score (Fig 5H). Other changes are seen within Sin3a-LOF/Sftpc-AT2 cells included a prominent induction of β6 integrin-Itgb6 (Fig 5I, J), which is involved in TGFβ1 activation (78, 79). In addition, immunofluorescence staining showed that β6+ AT2 cells accumulated proximal to fibrotic foci in Sin3a-LOF mice (Fig 5K, Supplemental Figure 9E). Collectively, these data demonstrate that TGFβ signaling is activated in AT2 epithelial cells of both human IPF- and Sin3a-LOF lungs.

**Figure 5.**
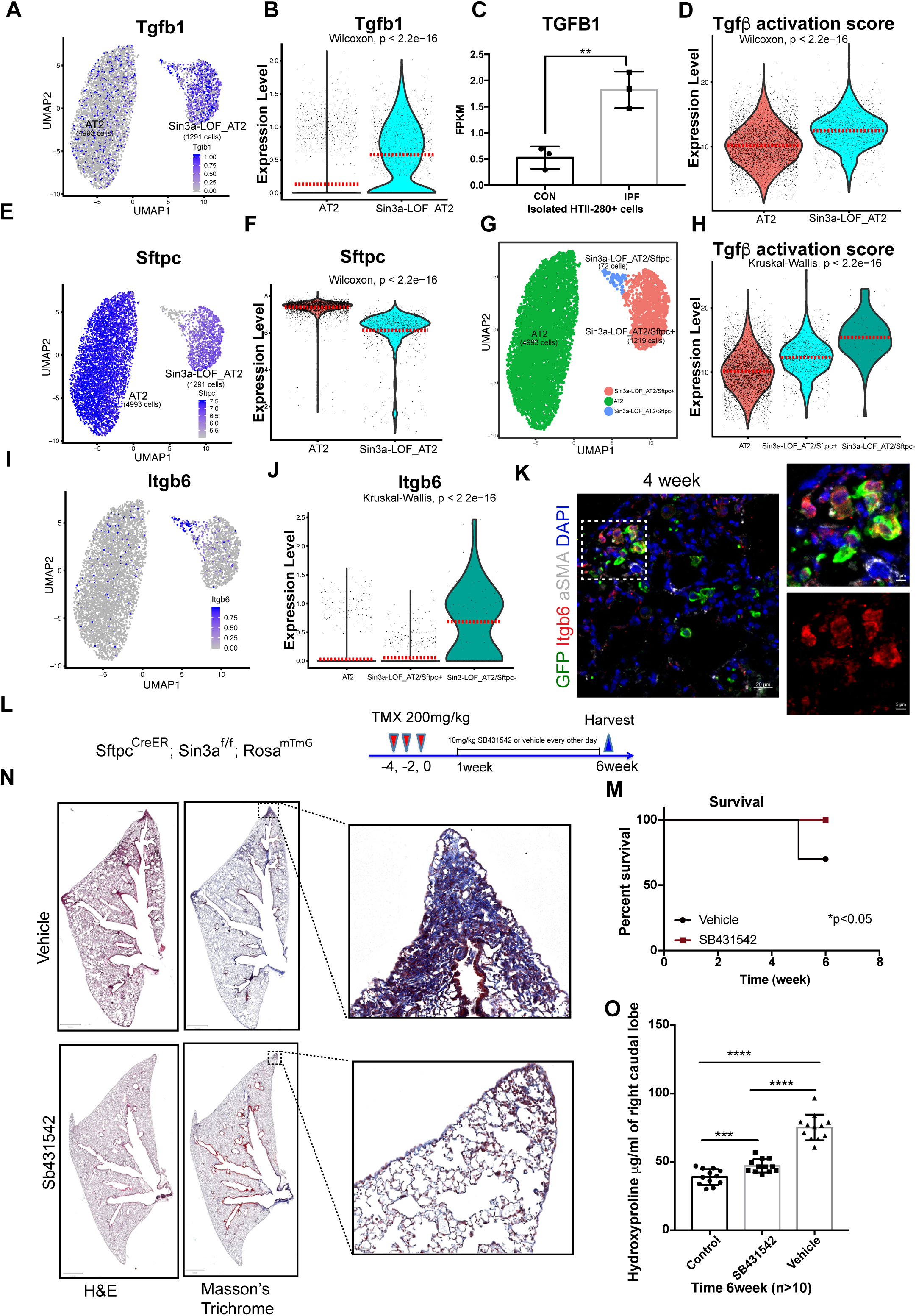
Inhibition of TGFβ mitigates lung fibrosis resulting from Sin3a-LOF in AT2 cells. A. UMAP visualization showing relative expression of Tgfb1 in mouse scRNA-Seq AT2 cell subsets. B. Violin plot representation showing relative Tgfb1 expression in mouse scRNA-Seq AT2 cells subset. Dash line represents median expression of each group. C. Normalized FPKM value of TGFB1 derived from bulk RNA-Seq of isolated human HTII-280 cells from control donor and IPF patient samples. D. Violin plot of Tgfβ activation score calculated using Tgfβ KEGG pathway genes in mouse scRNA-Seq AT2 cell subsets. E. UMAP visualization relative expression of Sftpc in mouse scRNA-Seq AT2 cell subsets. F. Violin plot representation of relative expression of Sftpc of mouse scRNA-Seq AT2 cells subset. Dash line indicates median expression of each group. G. UMAP visualization of mouse scRNA-Seq AT2 cell subset re-clustering base on expression of Sftpc. H. Violin plot of Tgfβ activation score of re-clustered AT2 cell subset. I. UMAP visualization showing expression of Itgb6 in mouse scRNA-Seq AT2 cell subsets. J. Violin plot representation showing relative expression of Itgb6 in re-clustered mouse scRNA-Seq AT2 cell subsets. K. Immunofluorescence staining of lineage reporter (GFP), Itgb6 and αSMA in Sin3a LOF mice 4 weeks post-tamoxifen exposure. L. Schematic outline of experiment design for Tgfβ inhibitor treatment. M. Survival curve of Sin3a-LOF mice comparing SB431542 treatment and vehicle treatment groups 6 weeks post-tamoxifen exposure. *p<0.05 by Log-rank (Mantel-Cox) survival analysis. N. Hematoxylin & Eosin (H&E) and Masson’s trichrome staining 6 weeks post-tamoxifen treatment comparing SB431542 and vehicle treatment groups. O. Hydroxyproline content in right caudal lobe of Sin3a-LOF mice treated with either SB431542 or vehicle, 6 weeks after tamoxifen treatment (n=12). p-value calculated by two-tailed student t-test. * p<0.05, ** p<0.01, *** p<0.001, **** p<0.0001.

To further test the role of altered TGFβ signaling as a mediator of progressive pulmonary fibrosis in Sin3a-LOF mice, we inhibited TGFβ pathway activation by systemic delivery of the activin receptor-like kinase-5 (Alk4/5) inhibitor SB431542 (80). 10-12-week-old Sin3a-LOF mice received SB431542 i.p. every other day after the first week of tamoxifen treatment (Fig 5L). We found that SB431542 treatment protected against mortality and body weight loss seen in vehicle control Sin3a-LOF mice (Supplemental Figure 9F, Fig 5M) and attenuated lung fibrosis, evident by reduction of fibrotic foci and collagen deposition (Fig 5N-5O). Further, we found that Sin3a-LOF AT2 cells expressed the senescence marker Cdkn1a/p21 after SB431542 treatment, suggesting that TGFβ activation and fibrogenesis are downstream of AT2 cell senescence (Supplemental Figure 9G). Taken together, we conclude that loss of Sin3a in AT2 cells results in TGFβ activation and that enhanced TGFβ signaling is a distal driver of progressive lung fibrosis seen following AT2 progenitor cell dysfunction in Sin3a-LOF mice.

### Senescence rather than loss of AT2 cells drives progressive lung fibrosis

To determine contributions of AT2 loss versus senescence in progressive fibrosis, we compared fibroproliferative responses seen in Sin3a-LOF mice with those in mice with direct ablation of AT2 cells achieved through conditional activation of diphtheria toxin A (Sftpc^CreER^; DTA^f/f^; Rosa^mTmG^, here on referred as DTA; Fig 6A). 10-12-week-old Sin3a-LOF and DTA mice were exposed to tamoxifen once every week for 8 weeks and evaluated at 4, 6, 8 and weeks after the first exposure (Fig 6A). Ablation efficiency was assessed by real-time quantitative PCR of Sftpc mRNA and by immunofluorescent staining for Sftpc (Supplemental Figure 10A, B). At the 8-week exposure time point, DTA mice experienced 10% mortality (2 out of 20 mice) with no significant decline in bodyweight (Fig 6B, C). In contrast, all Sin3a-LOF reached the euthanasia endpoint for bodyweight decline which was classified as mortality (Fig 6B, C). Upon histological analysis, both Sin3a-LOF and DTA showed tamoxifen-dependent parenchymal consolidation and matrix deposition by Masson’s trichrome staining at 4 weeks (Fig 6D). These data are consistent with a previous report showing that conditional expression of the diphtheria toxin receptor within AT2 cells promotes matrix deposition(81). However, despite histopathological evidence of progressive fibrosis seen in Sin3a-LOF mice, DTA mice recovered at 6- and 8-week exposure time points with no histological evidence of tissue remodeling and fibrosis (Fig 6D and E). Histological evidence of progressive fibrosis in Sin3a-LOF mice was matched by time-dependent increases in lung hydroxyproline content (Fig 6F) with no progressive increases in hydroxyproline content observed in lung tissue of DTA mice (Fig 6G). Taken together these data suggest that changes in the function and molecular phenotype of AT2 cells from Sin3a-LOF mice rather than simply loss of AT2 cells are responsible for progressive pulmonary fibrosis.

**Figure 6.**
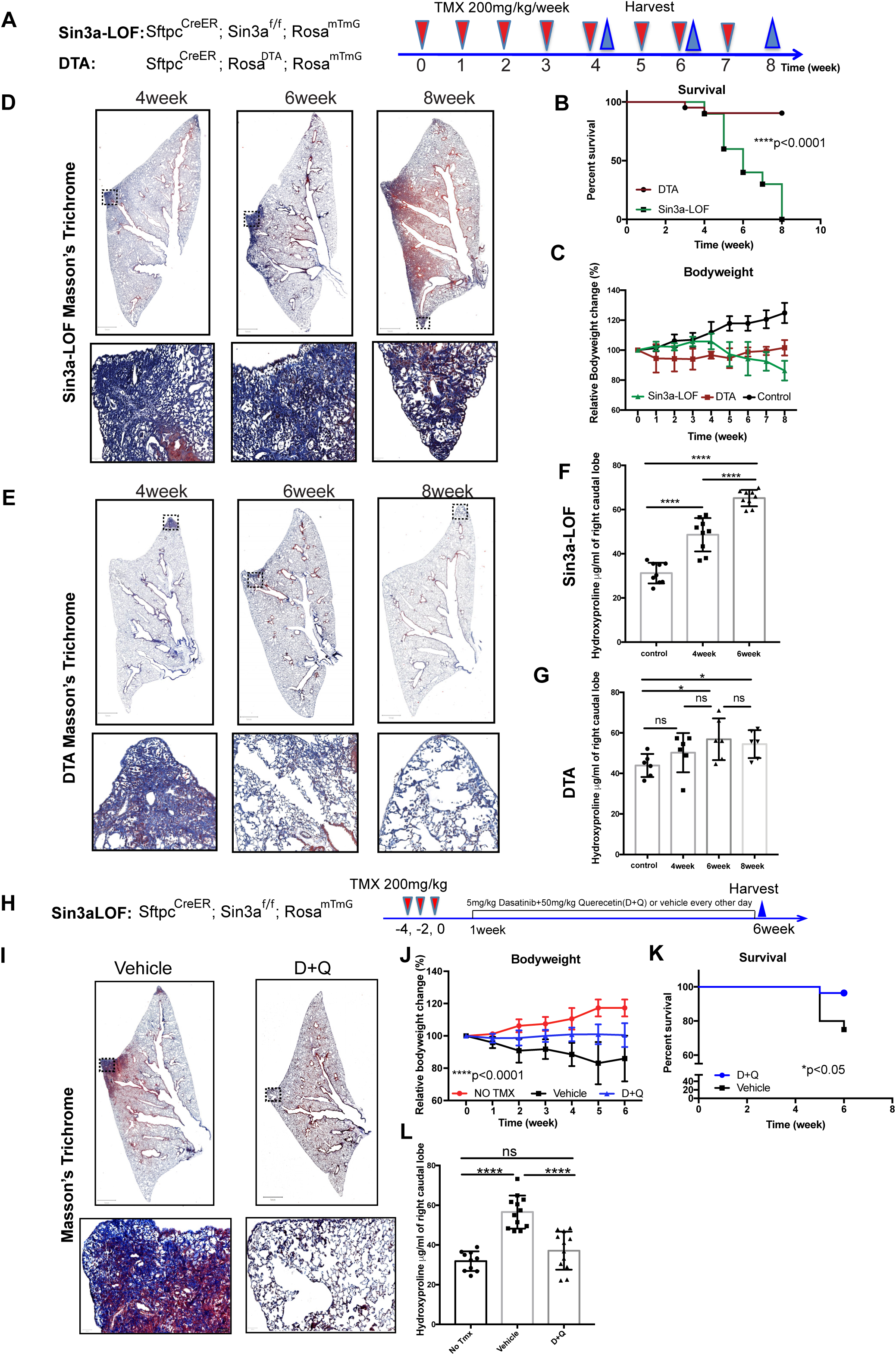
Senescence rather than loss of AT2 cells results in progressive lung fibrosis. A. Schematic outline of experiment design for AT2 cell ablation. B. Survival curve comparing Sin3a-LOF and DTA mice following repeated tamoxifen treatment. ****p<0.0001 by Log-rank (Mantel-Cox) survival analysis. C. Bodyweight change comparing Sin3a-LOF, DTA and control mice following repeated tamoxifen treatment. D. Masson’s trichrome staining of Sin3a-LOF mice following repeated tamoxifen treatment. E. Masson’s trichrome staining of DTA mice following repeated tamoxifen treatment. F. Hydroxyproline content in right caudal lobe of Sin3a-LOF mice following repeated tamoxifen treatment for 4, 6 repeated exposure of tamoxifen and control groups. G. Hydroxyproline content in right caudal lobe of DTA mice following repeated tamoxifen treatment for 4, 6, 8 repeated exposure of tamoxifen and control groups. H. Schematic outline of experiment design for senolytic drug treatment. I. Masson’s trichrome staining of Sin3a-LOF mice6 weeks post-tamoxifen treatment comparing 5mg/kg Dasatinib and 50mg/kg Quecertin (D+Q) cocktail treatment and vehicle treatment groups. J. Bodyweight change of Sin3a-LOF mice comparing D+Q treatment, vehicle treatment and control groups 6 weeks post-tamoxifen exposure. p<0.0001 for two-way ANOVA test and two-tailed student t-test comparing Vehicle vs D+Q treated group at each time point 2 weeks after tamoxifen exposure. K. Survival curve of Sin3a-LOF mice comparing D+Q and vehicle treatment groups 6 weeks post-tamoxifen exposure. *p<0.05 by Log-rank (Mantel-Cox) survival analysis. L. Hydroxyproline content in right caudal lobe of Sin3a-LOF mice treated with either D+Q or vehicle, 6 weeks after tamoxifen treatment (n>10 for each group). p-value calculated by two-tailed student t-test. * p<0.05, ** p<0.01, *** p<0.001, **** p<0.0001.

Finally, we tested the hypothesis that senescence of AT2 cells is the proximal driver of progressive pulmonary fibrosis observed following tamoxifen exposure of Sin3a-LOF mice. We treated 10-12-week-old Sin3a-LOF mice with a cocktail of senolytic drugs, Dasatinib and Quercetin (D+Q) or vehicle by oral gavage, every other day starting 1 week after the last dose of tamoxifen treatment for a total of 6 weeks. D+Q treatment protected Sin3a-LOF mice against bodyweight decline and mortality, in addition to mitigating histological evidence of fibrosis and expression of Cdkn1a/p21 (Fig 6I-K, Supplemental Figure 10C). Reduced fibrosis following D+Q treatment of Sin3a-LOF mice was also evident by a decrease in lung hydroxyproline content compared to vehicle treated Sin3a-LOF mice. Thus, we conclude that senescence of AT2 cells represents the proximal driver for progressive fibrosis seen in Sin3a-LOF mice and we speculate is a primary driver of tissue remodeling in IPF.

## Discussion

Even though fibrosis of epithelial tissues has a clear age-dependence, mechanisms by which advancing age contribute to initiation and/or progression of disease remain largely unknown. Increased cellular senescence is pathognomonic of disease and has been linked to both disease-associated gene variants and age-dependent loss of epithelial progenitor cell function. However, mechanisms by which cellular senescence drive tissue fibrosis are unclear. Herein we determine that senescence and declining abundance of alveolar type 2 (AT2) cells, stem cells that maintain the epithelial lining in the gas-exchange regions of the lung, are linked processes in end-stage idiopathic pulmonary fibrosis (IPF). In order to define disease mechanisms, we sought to define the contribution of stem cell senescence and stem cell loss towards initiation and progression of disease. We found that conditional loss of Sin3a within AT2 cells of adult mice induced a program of senescence and loss of progenitor cell function. Furthermore, even though either senescence or depletion of AT2 cells led to spontaneous lung fibrosis, fibrosis seen in mice with AT2 depletion was transient and rapidly resolved whereas conditional senescence promoted progressive interstitial lung fibrosis resembling that seen in human IPF. We establish that senescence, rather than stem cell depletion per se, represents a critical determinant of fibrogenesis in epithelial tissues.

Cellular senescence was first characterized as a state of cell cycle arrest after extensive proliferation and hence referred to as replicative senescence (RS). However, senescence can also be induced by multiple intrinsic stress-related stimuli such as telomere loss, DNA damage, oxidative stress, endoplasmic reticulum stress and oncogene activation (13, 82). These triggers induce cell cycle arrest through one of two major mechanisms, activation of either p53–p21 and/or p16INK4a–pRB pathways (72, 83, 84). We show that p53 signaling is one of the top pathways upregulated in AT2 isolated from IPF patient lung samples compared to their counterparts isolated from control donor lung tissue. Increased expression of CDKN1A/p21, a downstream target of p53, suggested that AT2 cell cycle arrest and senescence may depend on activation of the p53-p21 signaling axis. Certainly, hypertrophic AT2 cells present within IPF explant tissue show strong induction of p21 but also show elevated immunoreactivity for p16. These AT2 cells also show reduced immunofluorescent staining for the HDAC co-stimulatory factor Sin3a, and we were able to show that Sin3a LOF in mouse AT2 cells resulted in p53-p21 activation and p53-dependent senescence and fibrosis. Even though we cannot exclude a role for p16 in AT2 senescence, our data suggest that p53 activation is necessary (and sufficient) to induce AT2 senescence leading to progressive pulmonary fibrosis.

The senescence-associated secretory phenotype (SASP) has been shown to promote cell proliferation (85-87), stimulate cell motility (invasion, migration) (88-90), regulate cell differentiation (32, 88-90), and affect leukocyte infiltration (91-93), all of which serve important roles in repair following acute injury yet is a likely contributor to IPF pathogenesis. We show that a SASP gene expression signature is present within AT2 cells of the IPF lung and is induced following Sin3a loss in mouse AT2 cells. The critical role for AT2 senescence in both initiation and progression of fibrosis was suggested by progressive fibrosis of Sin3a-LOF mice but not with AT2 cell ablation and the demonstration that senolytic drugs were protective. Interestingly, a recent publication of a small Phase 1 study in IPF patients using the senolytic agents Dasatinib and Quercetin was performed and although the primary endpoint of change in Iung function was negative, patients reported improvements in breathlessness(94). Our finding that elimination of senescent AT2 cells from lungs of Sin3a-LOF mice protects from progressive fibrosis supports a critical role for AT2 senescence in both initiation and progression of IPF.

Both p53 and TGFβ were key upstream regulators in both IPF and Sin3a-LOF AT2 cells. Furthermore, we found that integrin alpha(v)beta6 (Itgb6), responsible for stretch-activation of latent TGFβ (79, 95), localized immediately adjacent to sites of lung fibrosis. Matrix deposition and fibroproliferation were inhibited by systemic delivery of the Alk4/5 inhibitor SB431542, indicating that TGFβ activation is an obligate downstream mediator of AT2 dysfunction. Given that TGFβ is a known regulator of fibrosis in many organs including the lung, our own data further highlight the importance of senescent AT2 cells as a source of TGFβ and, consistent with the previously reported work of Tatler and Jenkins (78, 79), that epithelial activation of TGFβ functions as a driving force for fibroproliferation. We therefore establish in both human IPF and in the Sin3a-LOF mouse model of IPF, that p53 activation and induction of SASP are proximal events leading to TGFβ activation and progressive fibrosis. Sin3a-LOF mice therefore serve as an animal model that recapitulates all pathologic hallmarks of IPF.

In summary, we present a novel conditional mouse model of spontaneous progressive interstitial lung fibrosis in which loss of Sin3a function and resulting senescence of AT2 cells is sufficient to bypass age-dependent changes to AT2 cells of the human lung that lead to IPF. Conditional Sin3a-LOF in mouse AT2 cells activates senescence in p53 and p21 dependent manner, leading to TGFβ activation, downstream signaling and fibrogenesis. Using this mouse model, we provide evidence that early targeting of senescent cells is of therapeutic benefit in moderating lung fibrosis, a strategy that may have broad potential to mitigate fibrosis in a wide range of epithelial tissues.

## Methods

### Study population

Explant tissue was obtained from patients undergoing transplantation for end-stage IPF at either University of California Los Angeles or Cedars-Sinai Medical Center, in compliance with consent procedures accepted by the Internal Review Board of Cedars-Sinai Medical Center. Human lung specimens obtained through the International Institute for the Advancement of Medicine (IIAM) were obtained in compliance with consent procedures developed by IIAM and approved by the Cedars-Sinai Medical Center IRB. Patient demographics provided in Supplemental Table 1.

### Mouse strains

Sftpc^CreER^ mice (The Jackson Laboratory, stock number 028054), Rosa26^mT/mG^ (The Jackson Laboratory, stock number 007576), RosaDTA mice (The Jackson Laboratory, stock number 009669), p53^flox/flox^ mice (The Jackson Laboratory, stock number 008462) were from The Jackson Laboratory. Sin3a^flox/flox^ mice were in previous publication Dannenberg et al., 2005 (96). All mice were maintained, and treatments were carried out according to Institutional Animal Care and Use Committee-approved protocols. 10- to 12-week-old mice are used for experiments in this study. For tamoxifen treatment, 200mg/kg of tamoxifen dissolved in corn oil was administered by intraperitoneal injection either 3 doses (every other day) or one dose/week to DTA and Sin3a-LOF comparison experiment due to acute death of DTA mice for the 3 doses of 200 mg/kg every other day treatment. All mice were closely observed for bodyweight and health, any mice reach to 20% bodyweight or more reduction will be euthanized and collected per institution animal regulation rules.

### Bleomycin injury

For bleomycin lung injury experiment, C57BL/6 wild-type (WT)mice 10- to 12-week-old were anesthetized by isoflurane inhalation, and bleomycin (1.25 units/kg, APP Pharmaceuticals, Schaumburg, IL) or sterile DPBS was intratracheally administered once. Lung collections are done at different time points (7, 14, 21).

### Drug treatment

SB431542 (Selleck Chemicals, S1067) was administered by intraperitoneal injection at a dose of 10mg/kg or vehicle control every other day 1 week after tamoxifen treatment until collection at 6 weeks post-tamoxifen treatment time point. Pifthrin-α (Selleck Chemicals, S2929) was administered by intraperitoneal injection at a dose of 3mg/kg or vehicle control every other day 1 week after tamoxifen treatment until collection at 6 weeks post-tamoxifen treatment time point. Dasatinib (Sigma Aldrich, CDS023389) and Quercetin (Sigma Aldrich, Q4951) were administered by intraperitoneal injection at a dose of 5mg/kg and 50mg/kg respectively in cocktail or vehicle control every other day 1 week after tamoxifen treatment until collection at 6 weeks post-tamoxifen treatment time point.

### Immunofluorescence staining, histology staining, immunohistochemistry, imaging and quantification

Immunofluorescence staining, Hematoxylin and Eosin (H&E) and Masson-Trichrome staining was performed on fixed lung tissue embedded in paraffin and processed as previously described (97). TSA amplifications (Perkin Elmer, NEL744001KT) per manufacture instruction are done for the staining of CDKN1A and CDKN2A. All immunofluorescence images were taken using a Zeiss 780 confocal microscope. Quantification is done either using FIJI or Imaris. Proteins were quantitated using bicinchoninic acid (BCA) protein assay (Pierce/Thermo Scientific, 23225) per manufacturer’s instruction. Western blot and hydroxyproline assays were performed as previously published (98). Noted that hydroxyproline assays were performed either right lobes (Figure 3) or right caudal lobe for the rest.

Primary antibodies used for immunofluorescence staining included:chicken anti-GFP (1:1000, Abcam, ab13970), rabbit anti-proSPC (1:1000, Millipore, AB3786), mouse anti-smooth muscle actin (1:10,000, Sigma-Aldrich, A2547), rabbit anti-Sin3a (1:100, Santa Cruz Biotechnologies, sc-767), mouse anti-CDKN1A (1:100, Santa Cruz Biotechnologies, sc-6246), mouse-HTII-280 (IgM) (1:500, Terrace Biotech, TB 27AHT2-280), mouse anti-CDKN2A (IgG2b) (1:100, Abcam, ab54210), rabbit anti-CDKN2A(1:100, Abcam, ab3642), rabbit anti-Gapdh (1:1000, Cell signaling Technology, 2118).

Secondary antibodies were: Alexa Fluor 647-conjugated donkey anti-mouse IgG(H+L) (1:1000, Life Technologies, A31571), Alexa Fluor 594-conjugated donkey anti-rabbit IgG(H+L) (1:1000, Life Technologies, A21207), Alexa Fluor 488-conjugated donkey anti-rabbit IgG(H+L) (1:1000, Life Technologies, A21206), Alexa Fluor 488-conjugated donkey anti-mouse IgG(H+L) (1:1000, Life Technologies, A21202), Alexa Fluor 488-conjugated goat anti-chicken IgG(H+L) (1:1000, Life Technologies, A11039) and Alexa Fluor 488-conjugated donkey anti-chicken IgY (IgG) (H+L) (1:1000), FITC Goat anti-mouse IgM (1:1000, ThermoFisher Scientific, 31992), Alexa Fluor 568-conjugated goat anti-mouse IgG2b (1:1000, ThermoFisher Scientific, A-21144), goat anti-rabbit HRP-conjugated (1:3000, Jackson ImmunoResarch Laboratories, 111-0350144), goat anti-mouse HRP-conjugated (1:3000, Jackson ImmunoResarch Laboratories, 111-035-003).

### Quantitative real-time PCR

Complementary DNA was synthesized from total RNA by using iScript Reverse Transcription Supermix (Bio-Rad). Quantitative RT-PCR was performed using the SYBR Green Master Mix (Bio-Rad). Gapdh expression values were used to control for RNA quality and quantity. Experiments were performed with at least three independent biological replicates and data shown represent the average ±SEM.

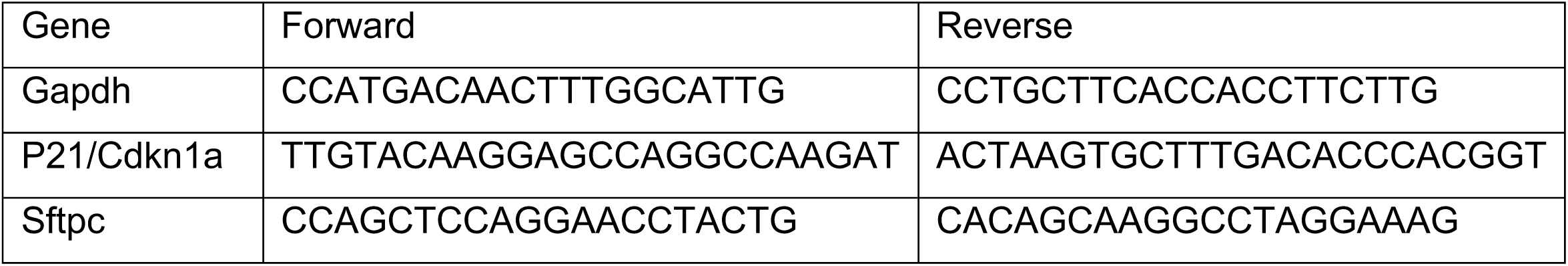

### Senescence-associated β-galactosidase detection

Both mouse and human lung samples were fixed 10% formalin for 15 minutes for further frozen sections. Lung sections are stained using Senescence Cells Histochemical Staining Kit (Sigma-Aldrich, CS0030-1KT) to detect senescence-associated β-galactosidase detection following the manufacturer’s recommendations. Tiled images were taken using Zeiss Axio observer Z1 microscope with color camera.

### Cell isolation

Mouse epithelial isolation was processed as described previously with the following modifications(73). After mouse lung tissue was instilled enzymatic digestion with elastase (4 U/ml) (Worthinton Biotechnical, LS002280), mouse lung tissues were diced into small pieces and further digested with 1x Liberase (0.25U/ml) (Sigma-Aldrich, 5401127001) on a Thermomixer at 37°C for 30 min. Dissociated single-cell preparations were stained for enriching for epithelial cells and depleting of leukocytes, and endothelial cells using antibodies against the following molecules: Epcam, CD45, and CD31 (Biolegend). BD Influx cell sorter (Becton Dickinson) was used to fluorescence-activated cell sorting (FACS) enrich epithelial cells for further scRNA-Seq. Viability was determined by staining cell preparations with DAPI (ThermoFisher Scientific), 15 minutes prior to cell sorting. AT2 cells enrichment will further gating for fluorescent report GFP (DAPI-,CD31-, CD45-, Epcam+,GFP+).

Human IPF explant and donor tissues were processed as described previously with the following modifications(20). IPF fibrotic region was identified by pathologists at Biobank at Cedars-Sinai Medical Center and UCLA, further validated by H&E staining. Lung tissue was finely minced and washed in Hank’s balanced salt solution (HBSS) (Corning) at 4°C for 5 minutes with rocking, followed by centrifugation for 5 minutes at 600g and 4°C. Repeat as necessary to remove as much blood as possible. The minced cleaned tissue was then incubated in HBSS containing 1X Liberase (0.25U/ml) (Sigma-Aldrich, 5401127001) and DNaseI (0.5mg/ml) (Qiagen, 1023640), incubated with a Thermomixer (Eppendorf) and agitate at 1000 rpm at 37°C for 30 min. Tissue chunks will be further dissociated by going through 16 G syringe needles. Agitate gently at 1000 rpm at 37°C on thermomixer for another 15 min. Dissociated single-cell preparations were stained for enriching for epithelial cells and depleting of erythrocytes, leukocytes, and endothelial cells using antibodies against the following molecules: EPCAM, CD235a, CD45, and CD31 (Biolegend). BD Influx cell sorter (Becton Dickinson) was used to fluorescence-activated cell sorting (FACS) enrich epithelial cells for further scRNA-Seq. Viability was determined by staining cell preparations with DAPI (ThermoFisher Scientific), 15 minutes prior to cell sorting.

### *In vitro* organoid cultures

Three thousand flow sorted DAPI-CD45-CD31-Epcam+GFP+ cells were cultured with one hundred thousand MLg mouse lung fibroblasts and Matrigel (BD Biosciences) seeded onto Transwell filter inserts (BD Biosciences) as previously described (97).

### Bulk RNA-seq

RNA was extracted from lung tissue of Sftpc^CreER^; Sin3a^flox/flox^; Rosa26^mT/mG^ mice (experimental group) at different time points, using miRNeasy Mini Kit (Qiagen, 217004); or AT2 cells that were isolated from Sftpc^CreER^; Sin3a^flox/flox^; Rosa26^mT/mG^ and Sftpc^CreER^; Rosa26^mT/mG^ 2 weeks after tamoxifen treatment using a miRNeasy Micro Kit (Qiagen, 217084). Library preparation and sequencing were performed by Cedars-Sinai Genomic Core using Illumina NextSeq 500 (Illumina) with single-end 75bp sequencing chemistry. On average, about 20 million reads were generated from each sample. Raw reads were aligned using Star aligner 2.6.0(99). Counts are determined by featureCounts 1.6.3(100). Differential gene expression was determined by DEseq2(101). Top 100 differential genes were determined based on fold change and test statistics. The whole analysis pipeline was run on Galaxy server (use.galaxy.org).

### scRNA-Seq

For mouse scRNA-Seq of Sin3a-LOF vs control, CD45-CD31-Epcam+ lung epithelial cells from adult lungs of 4 male and 4 female mice for each group were FACS sorted, pooled and performed for scRNA-seq using the 10x Genomic Chromium system. For bleomycin treated mice vs control, we pooled three mice for each group, and hashed cells from each mouse with mouse cell hashing antibodies (Totalseq™, BioLegend, San Diego, CA) before pooling and sorting. Viable lung cell suspension (DAPI^-^) was flow sorted for lung immune cells (CD45^+^), endothelial cells (CD31^+^), epithelial cells (CD326^+^) and stromal cells (CD45 ^-^CD31^-^CD326^-^). Equal proportions of all four cell types were pooled and subjected for single cell capture. Similarly, for human scRNA-Seq, CD45-CD31-EPCAM+ lung epithelial cells were FACS sorted from either IPF explant or donor samples and performed for scRNA-Seq using the 10x Genomic Chromium system. Single cells were captured using a 10X Chromium device (10X Genomics) and libraries were prepared according to the Single Cell 3’ v2 Reagent Kits User Guide (10X Genomics). The barcoded sequencing libraries were quantified by quantitative PCR using the KAPA Library Quantification Kit (KAPA Biosystems, Wilmington, MA). Sequencing libraries were sequenced using Novaseq 6000 (Illumina) with either custom sequencing setting of 26bp and 98bp for read 1 and 2 respectively, or 150 bp pair-end to obtain a sequencing depth of ∼5×10^4^ reads per cell.

### Data analysis

CellRanger v2.2.0 software was used with the default settings for de-multiplexing, aligning reads with STAR software to modified GRCM38/mm10 mouse reference genome (reporter gene GFP was added to the reference genome) for mouse scRNA-Seq, GRCh38 transcriptome reference genome for human scRNA-Seq, and counting unique molecular identifiers (UMIs). Cell hashing data were de-multiplexed using CITE-seq Count 1.4.1. Downstream analysis was performed using Seurat v2.3.4 R package (102). Canonical correlation analysis (CCA) was applied to integrate and combining data sets for unsupervised clustering. tSNE(t-distributed stochastic neighbor embedding) and UMAP (Uniform Manifold Approximation and Projection) method have been used for visualization of unsupervised clustering. The cell type of each cluster is determined by known markers of individual cell types. For scRNA-Seq data from Reyfman et al. (34), to show a balance cell number of AT2 cells from donor and IPF samples, we random subset 3000 AT2 cells from control group to analyze together with AT2 cells from donor samples.

### Data and code availability

All the sequencing data along with their associated metadata have been deposited in the GEO database under accession code GSE132908, GSE132909, GSE132910, GSE132914 and GSE131800. All relevant data and code of this study are available from the authors upon reasonable request. Note: part of mouse scRNAseq data (control data set also available through GSE131800)(103)

### Statistical analysis

One-way ANOVA analysis was used for multiple comparisons. Two-way ANOVA analysis was used for multiple group and factor comparisons (bodyweight changes along time points and between groups). Two-tailed, unpaired Student’s t-test (Prism, GraphPad) was used for two-group analysis. All analysis was performed in GraphPad Prism 7/8. Results were presented as the mean ± SEM. P<0.05 was considered statistically significant and significance levels are presented as *P<0.05, **P<0.01, ***P<0.001, ****P<0.0001.

## Author Contributions

Y, GC, and BRS contributed to conceptual design of experiments. CY, XG, TP, XL, GH performed experiments. CY and BRS analyzed data, HJS, SSW, DJ, PC, JAB, WCP, PWN, GD provided material and technical assistance. CY, PC and BRS were involved in manuscript writing.

## Acknowledgments

We especially thanks for Dr. William C Parks for suggestions on experimental design, critical reading and editing on the manuscript. We are grateful to the Cedars-Sinai Genomics Core for their assistance with bulk RNA-Seq and Biobank for assistance for tissue procuring. We also thank the Stripp lab members for suggestions and critical reading of the manuscript. We also thank for Bindu Konda, Guangzhu Zhang, Matthew Kostelny, Ningshan Liu, Vrishika Kulur for technical and material support. This study was funded by the California Institute of Regenerative Medicine (LA1-06915), the National Institutes of Health (4U01HL110967-05, P01HL108793 and T32HL134637), Celgene IDEAL Consortium and the Parker B. Francis Fellowship Program.

