## Supplemental Table 1 for "Senescence of alveolar stem cells drives progressive pulmonary fibrosis"

| Sample Name | Age | Gender | Smoking History |
| --- | --- | --- | --- |
| IPF1 | 74 | M | Never Smoker |
| IPF2 | 64 | F | Never Smoker |
| IPF3 | 68 | F | Never Smoker |
| IPF02-17 | 70 | M | Former Smoker (quit date 1/1/08) |
| IPF08-17 | 65 | M | Unknown |
| IPF09-17 | 75 | M | No Significant Smoking history |
| IPF10-17 | 63 | M | Former smoker (quit date: 1/1/97, 1.3 ppd, pack-years: 36.4) |
| IPF01-18 | 64 | F | Never Smoker |
| IPF02-18 | 62 | F | Packs/day:0.30, years 9.00, pack years:2.7 |
| IPF04-18 | 65 | F | Never Smoker |
| CON1 | 79 | F | Never Smoker |
| CON2 | 65 | F | Smoker |
| CON3 | 62 | F | Never Smoker |
| CC003-14 | 17 | M | Never Smoker |
| CC004-14 | unknown | unknown | unknown |
| CC005-14 | 22 | M | unknown |
| CC007-14 | 62 | M | 2 pack/day for 30 years |
| CC009-14 | 75 | M | 1 pack/day for 20 years |
| CC05-17 | 58 | M | 1 Cigarette/day for 15-20 years |
| CC02-18 | 17 | F | Never Smoker |
| CC05-18 | 63 | F | Cigarettes, ½ ppd x 20yrs |
| CC07-18 | 61 | M | Never Smoker |
