## Supplemental Figure for "Senescence of alveolar stem cells drives progressive pulmonary fibrosis"

#### Supplemental Figure 1.

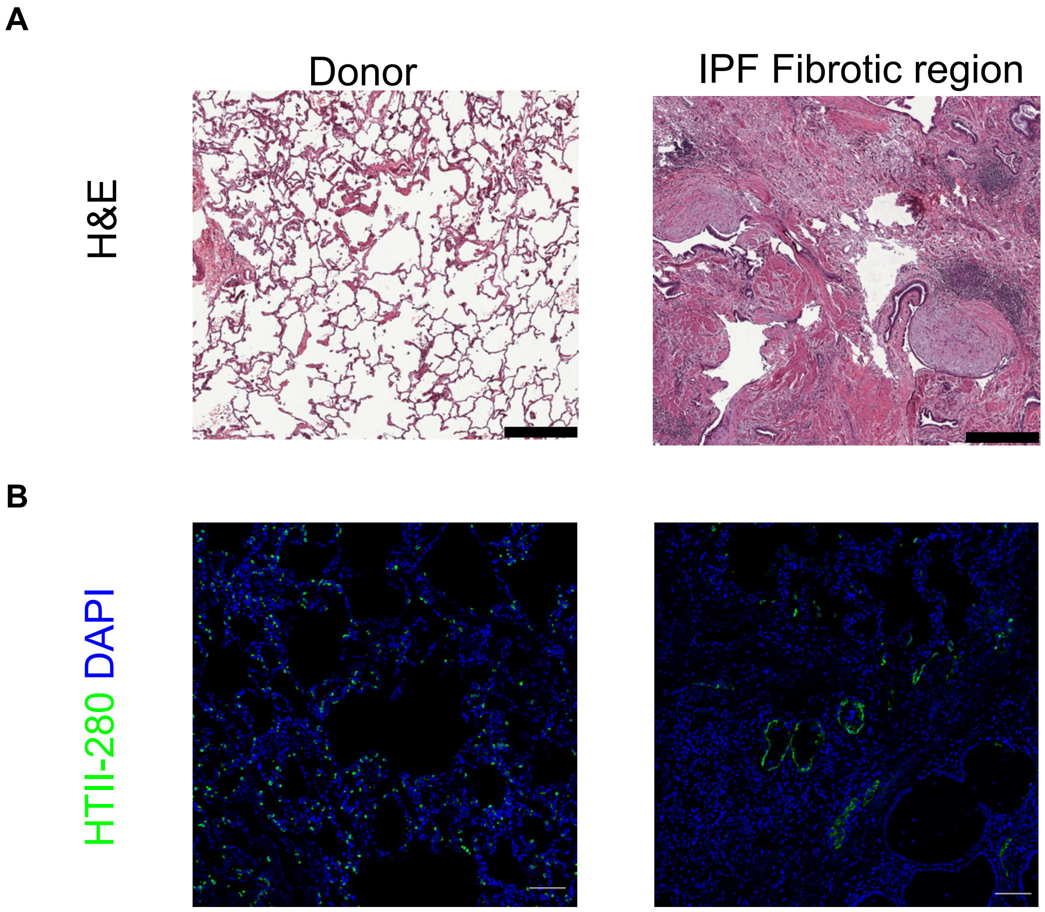


**Supplemental Figure1. Tissue remodeling in of fibrotic region of IPF patient lung tissues (Related to Fig 1).**
A. Representative image of H&E staining of IPF fibrotic region and donor distal lung tissue for scRNA-seq. Scale bar = 200μm. B. Representative immunofluorescence staining of HTII-280 of human control donor distal lung and IPF patient fibrotic lung tissue for scRNA-Seq. Scale bar = 50μm.

#### Supplemental Figure 2.

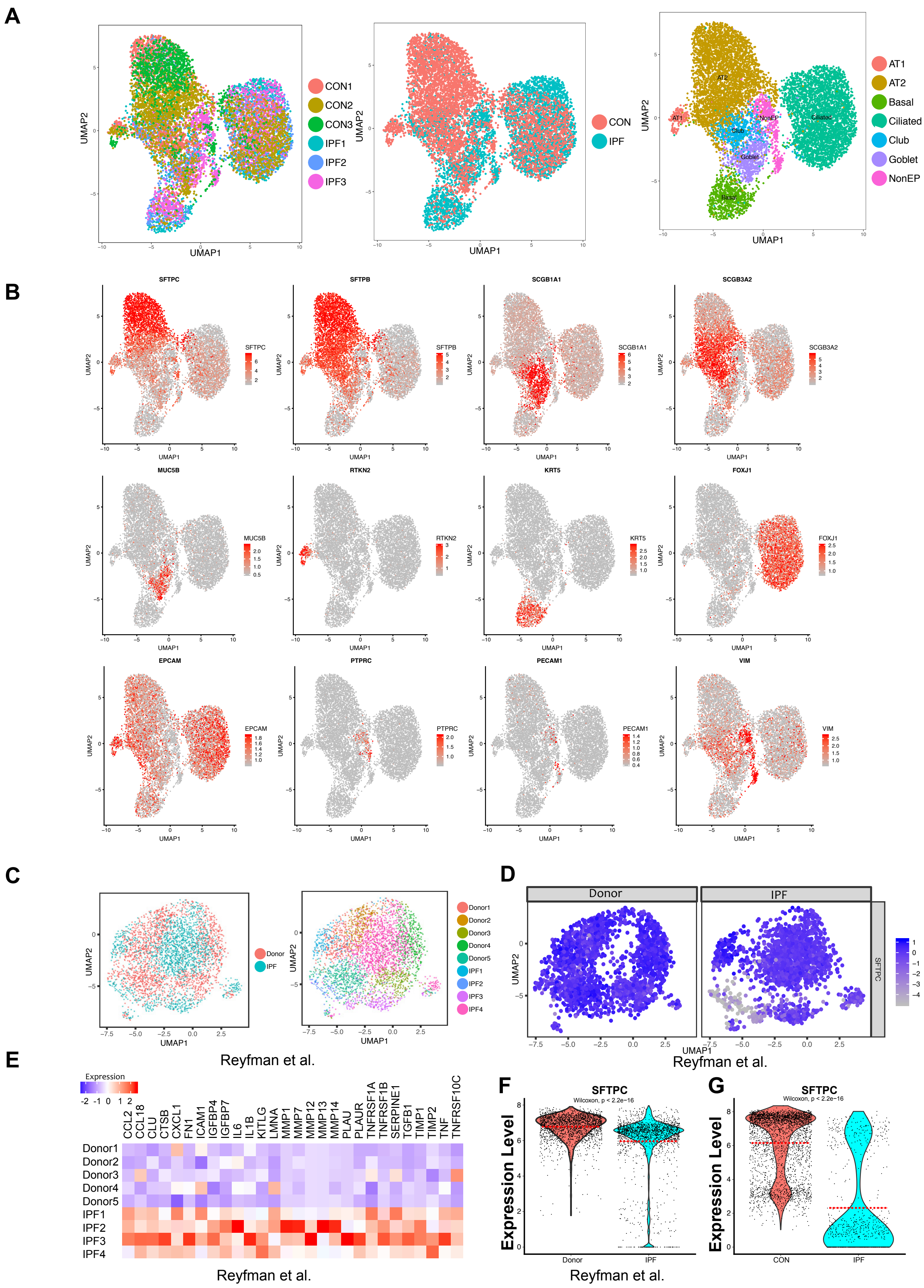

**Supplemental Figure 2. scRNA-Seq revealed senescent AT2 cells accumulation in fibrotic region of IPF patient lung (Related to Fig 1).**

A. UMAP visualization of cell origin, grouping and cell type clustering in human epithelial cells scRNA-Seq data on isolated epithelial cells from donor distal lung tissues and fibrotic region of IPF patient lung tissue. B. UMAP visualization of relative expression of known cell type specific markers used for cell type clustering. C. UMAP visualization of AT2 cell subsets, origin and clustering, from Reyfman et al. scRNA-Seq (random subset 3000 cells max per group). D. UMAP visualization showing relative expression of SFTPC comparing control and IPF from Reyfman et al. scRNA-Seq AT2 cell subsets. E. Heatmap of known SASP genes for average expression these genes of AT2 cells from each patient sample (AT2 cell subsets from Reyfman et al. scRNA-Seq). F. Violin plot representation showing relative expression of SFTPC in Reyfman AT2 cell subsets. Dash line indicates median expression of each group. G. Violin plot representation showing relative expression of SFTPC AT2 cell subsets from human epithelial cell scRNA-Seq. Dash line indicates median expression of each group.

#### Supplemental Figure 3.

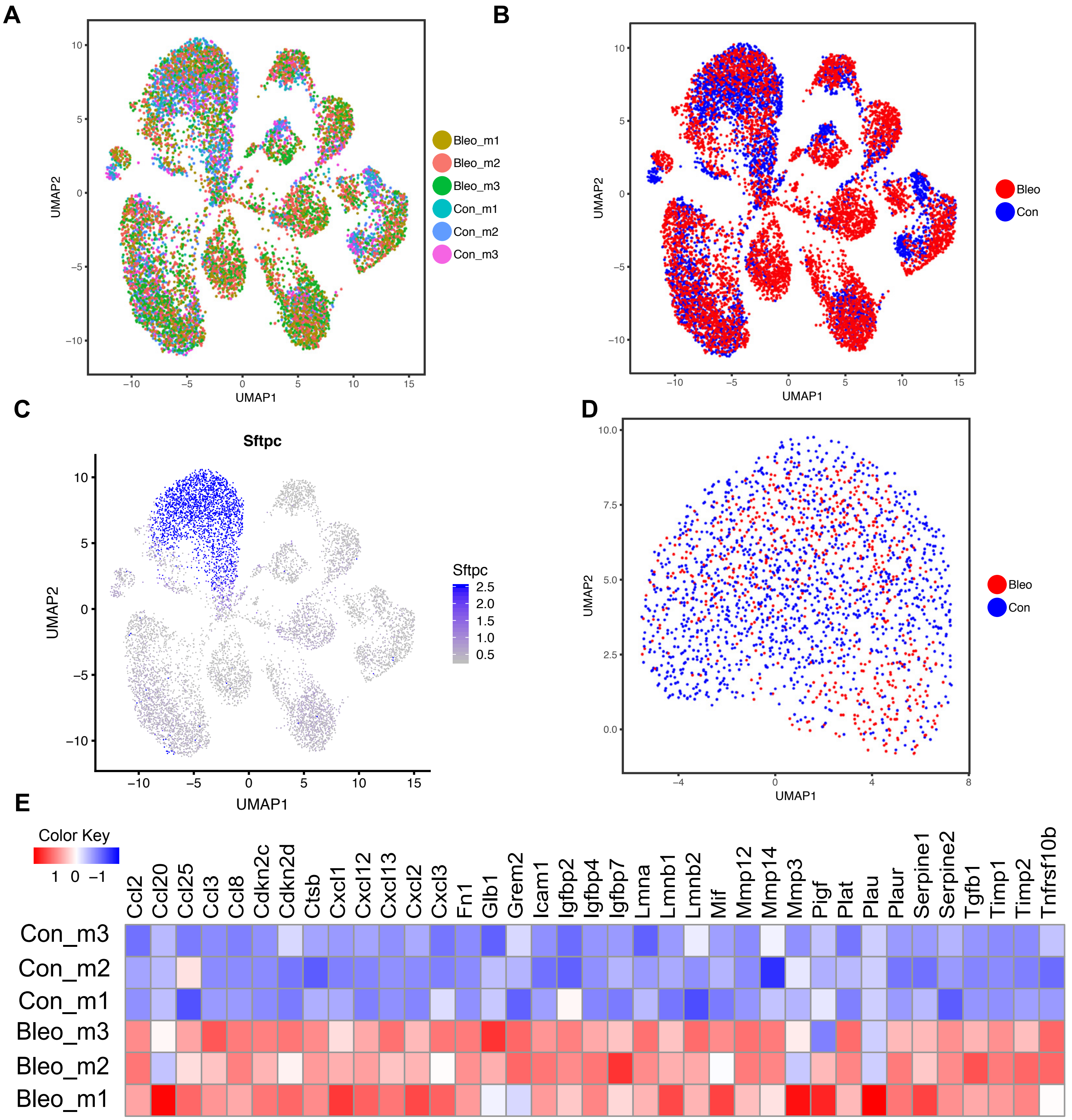


**Supplemental Figure 3. AT2 cell cellular senescence in Bleomycin treated mouse lung fibrosis model.**
A & B. UMAP visualization of cell origin and grouping from day 21 bleomycin treated lung samples and age matched saline treated mouse lung samples. B. UMAP visualization showing relative expression of SFTPC. D. UMAP visualization of AT2 cell subset. E. Heatmap of known SASP genes and for average expression of these genes in AT2 cells from each mouse sample.

### Supplemental Figure 4.

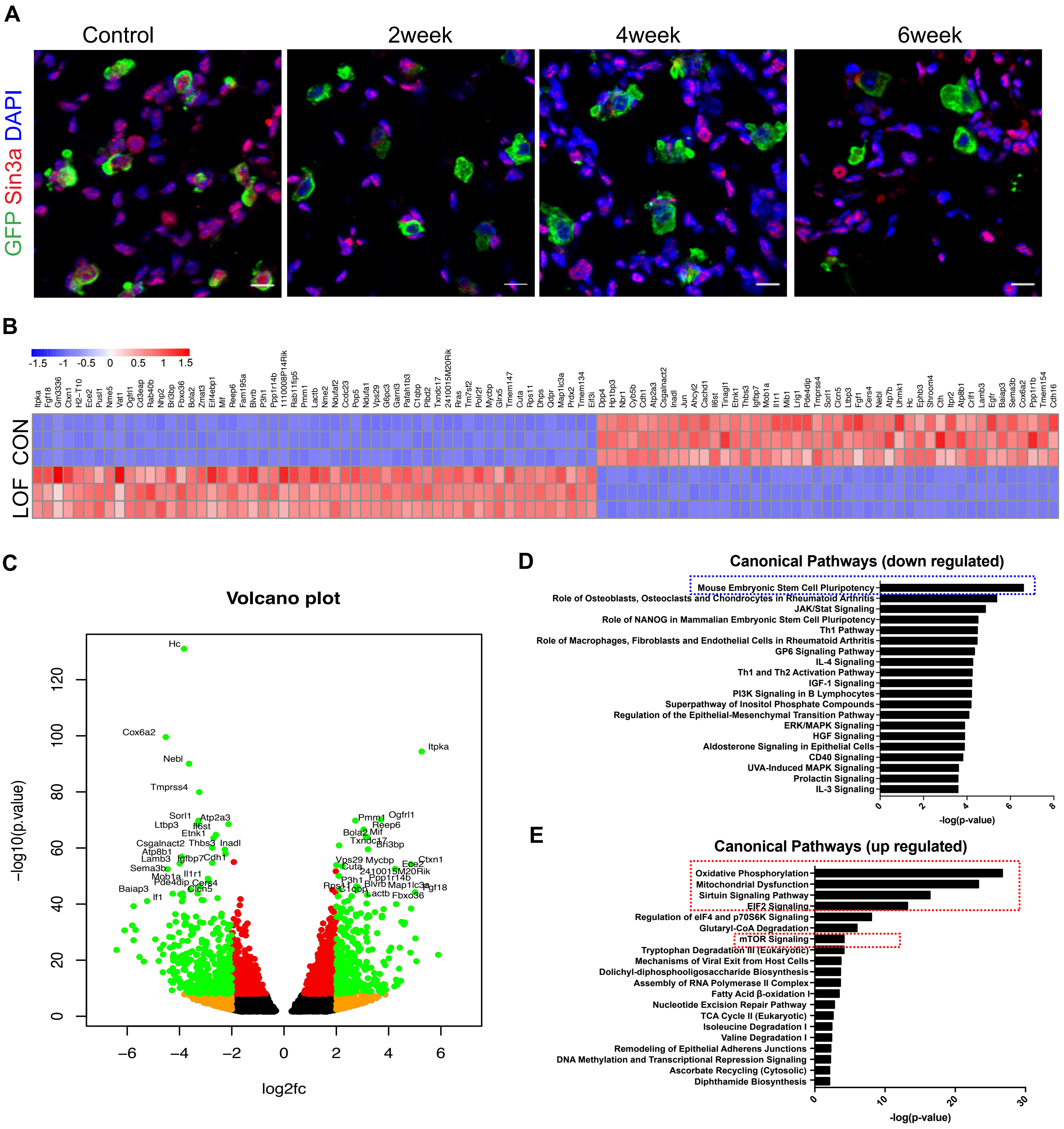


**Supplemental Figure 4. Bulk RNA-Seq reveals loss of Sin3a in adult AT2 cell leads to AT2 cell cellular senescence (Related Fig 2).**

A. Representative immunofluorescence staining of lineage reporter (GFP) and Sin3a of Sin3a-LOF mouse lung tissue samples over time course. Scale bar 20μm. B. Top 100 differential expression genes of RNA-Seq of isolated AT2 cells 2 weeks after tamoxifen treatment. C. Volcano Plot of differential expression genes of RNA-Seq of isolated AT2 cells 2 weeks after tamoxifen treatment. IPA Canonical pathway analysis of down-regulated genes D), up-regulated genes E) of Sin3a-LOF AT2 cells comparing control AT2 cells from RNA-Seq of isolated AT2 cells 2 weeks after tamoxifen treatment.

#### Supplemental Figure 5.

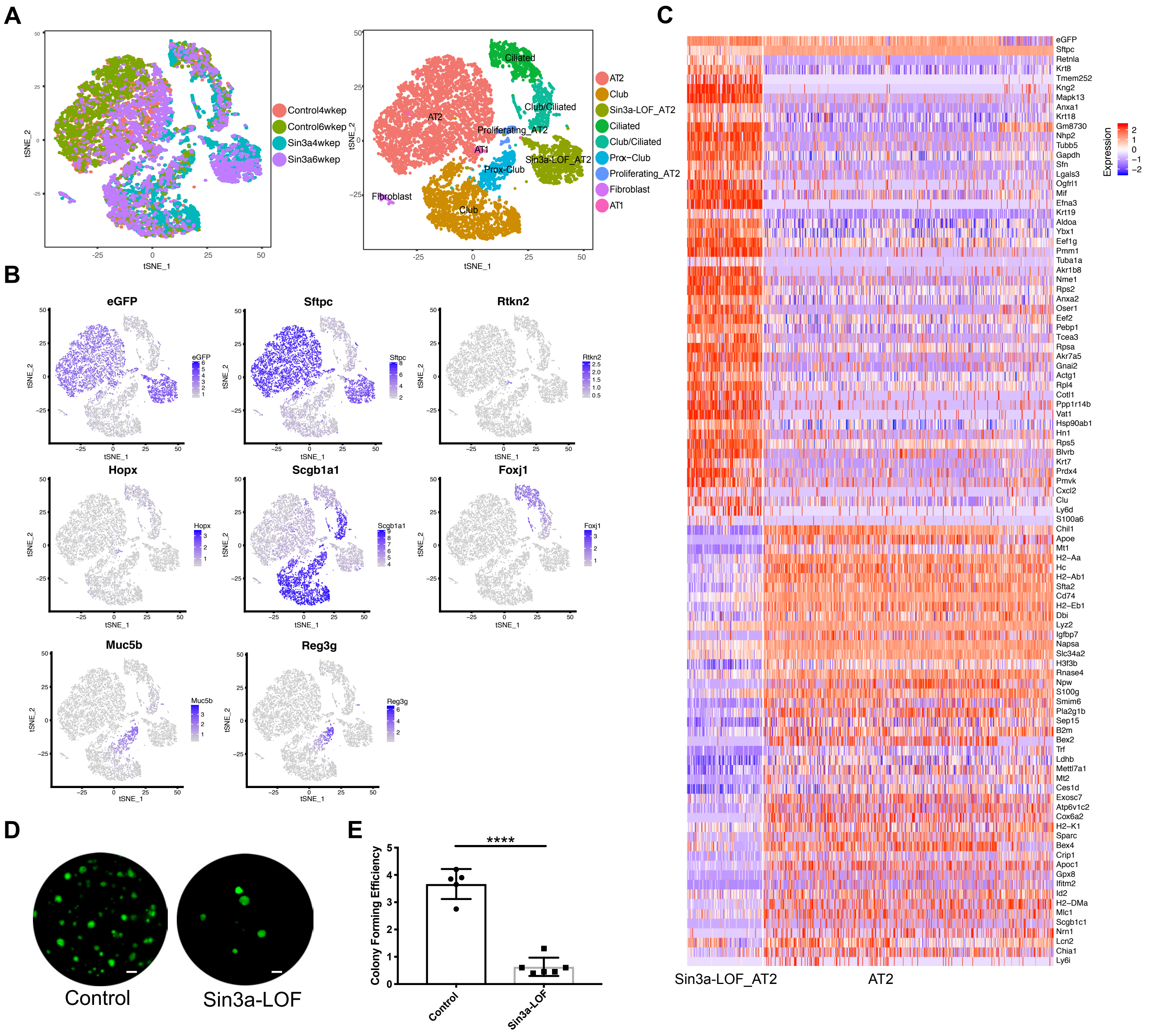


**Supplemental Figure 5. scRNA-Seq reveal loss of Sin3a in adult AT2 cell leads to AT2 cell cellular senescence (Related Fig 2).**
A. tSNE visualization of cell origin and cell type clustering for mouse epithelial cells scRNA-Seq isolated from 4 and 6 weeks after tamoxifen treatment of Sin3a-LOF lung samples and age matched control mouse lung samples. B. tSNE visualization of relative expression of known cell type specific markers used for cell type clustering. C. Heatmap of top 100 differential expression genes comparing Sin3a-LOF AT2 cells with control AT2 cells. D. Representative image of in vitro 3D organoid culture of Sin3a-LOF AT2 cells and control AT2 cells. Scale bar= 500µm. E. Colony forming efficiency quantification of 3D organoid culture. p-value calculated by two-tailed student t-test. * $p<0.05$, ** $p<0.01$, *** $p<0.001$, **** $p<0.0001$.

### Supplemental Figure 6.

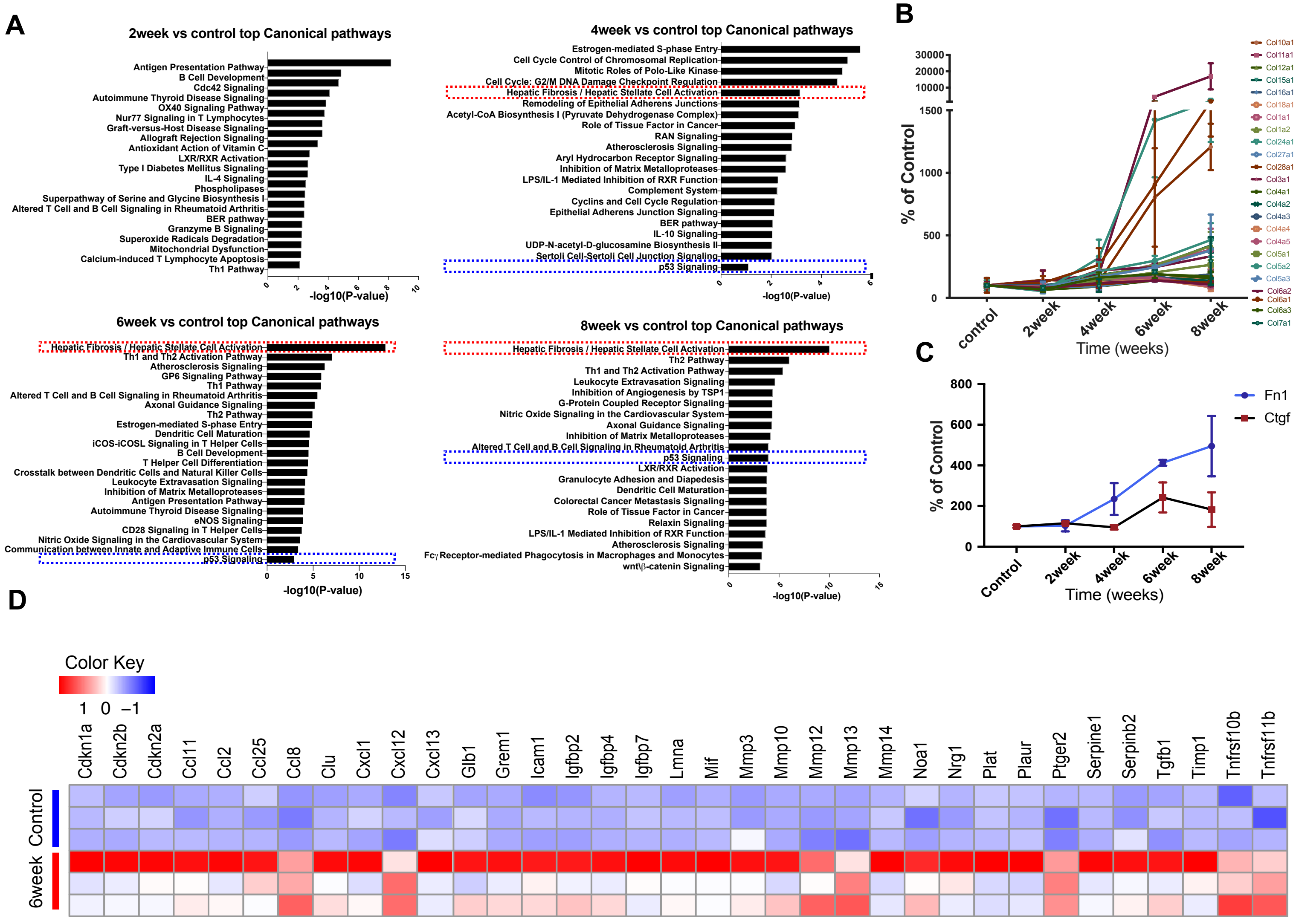


**Supplemental Figure 6. AT2 cell cellular senescence results in progressive lung fibrosis (Related to Fig 3 and 4).**

A. IPA Canonical pathway analysis of total lung tissue time course RNA-Seq. Fibrosis pathways are highlighted in red, p53 signaling pathways are highlighted in blue. B. Relative expression of collagens derived from total lung time course RNA-Seq. C. Relative expression of Fn1 and Ctgf derived from total lung time course RNA-Seq. D. Heatmap of SASP genes expression Sin3a-LOF lung 6 weeks post-tamoxifen treatment with treatment control derived from total lung time course RNA-Seq.

### Supplemental Figure 7.

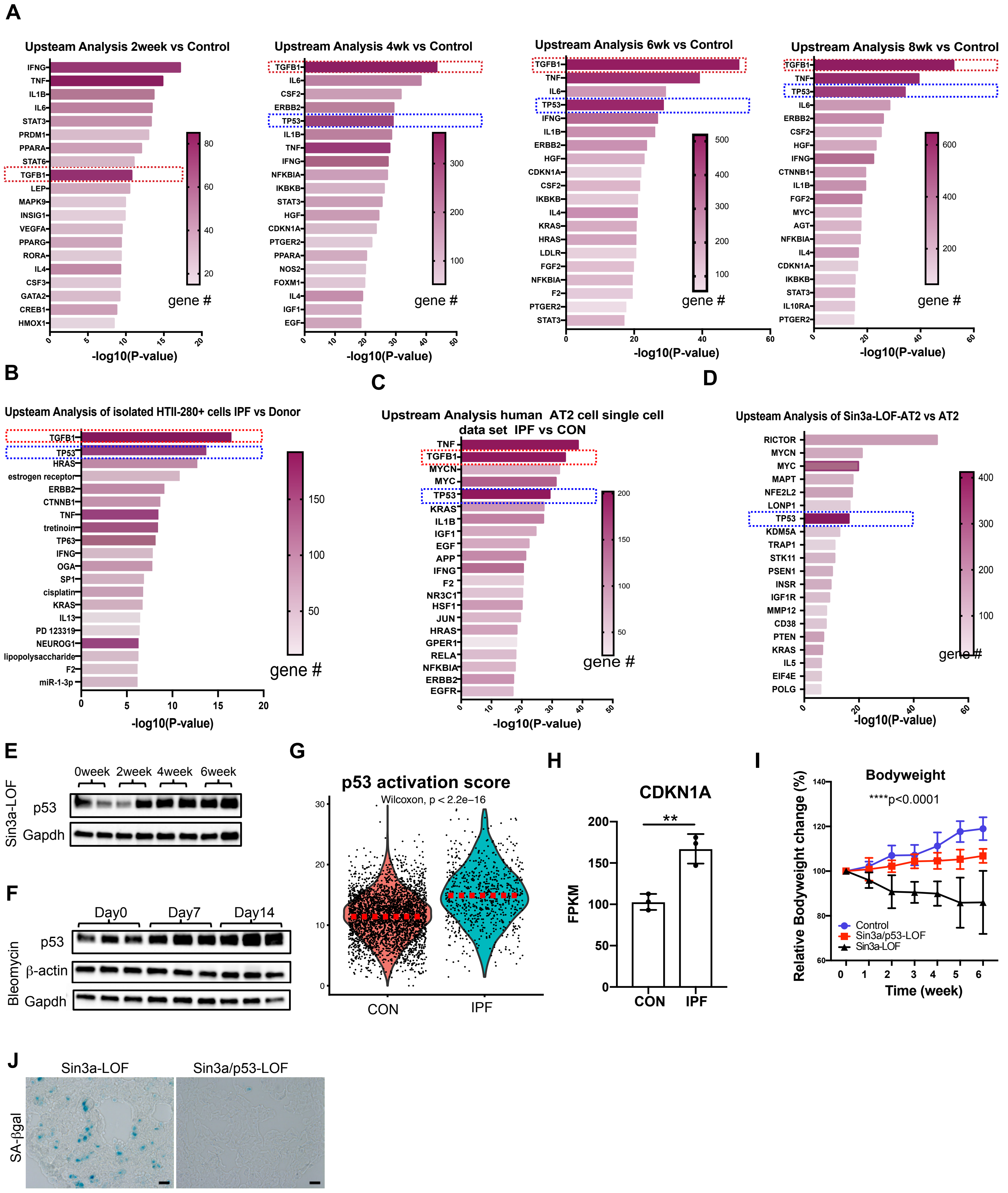

**Supplemental Figure 7. p53 signaling pathway activation in Sin3a-LOF AT2 cell cellular senescence (Related Fig 4 and Fig 5)**.

A. IPA upstream analysis of Sin3a-LOF total lung time course RNA-Seq. TGFB1 is highlighted in red, TP53 is highlighted in blue. B. IPA upstream analysis of bulk RNA-Seq of isolated human HTII-280 cells. TGFB1 is highlighted in red, TP53 is highlighted in blue. C. IPA upstream analysis of AT2 cell subsets from human epithelial cell scRNA-Seq. TGFB1 is highlighted in red, TP53 is highlighted in blue. D. IPA upstream analysis comparing Sin3a-LOF AT2 cells with control AT2 cells from mouse epithelial scRNA-Seq. TP53 is highlighted in blue. E. Western Blot detection p53 abundance within lung tissue of Sin3a-LOF mice. F. Western Blot detection p53 abundance within lung tissue of bleomycin treated mice. G. Violin plot representation showing p53 activation score of AT2 cell subset human epithelial cell scRNA-Seq. H. CDKN1A transcript level derived from bulk RNA-Seq of isolated human HTII-280 cells from control donor and IPF patient samples. I. Bodyweight change for Control, Sin3a-LOF and Sin3a/p53-LOF groups 6 weeks post-tamoxifen treatment. ****$p<0.0001$. J. SA-βgal signaling of Sin3a-LOF and Sin3a/p53-LOF mouse lung (6 weeks post-tamoxifen).

#### Supplemental Figure 8.

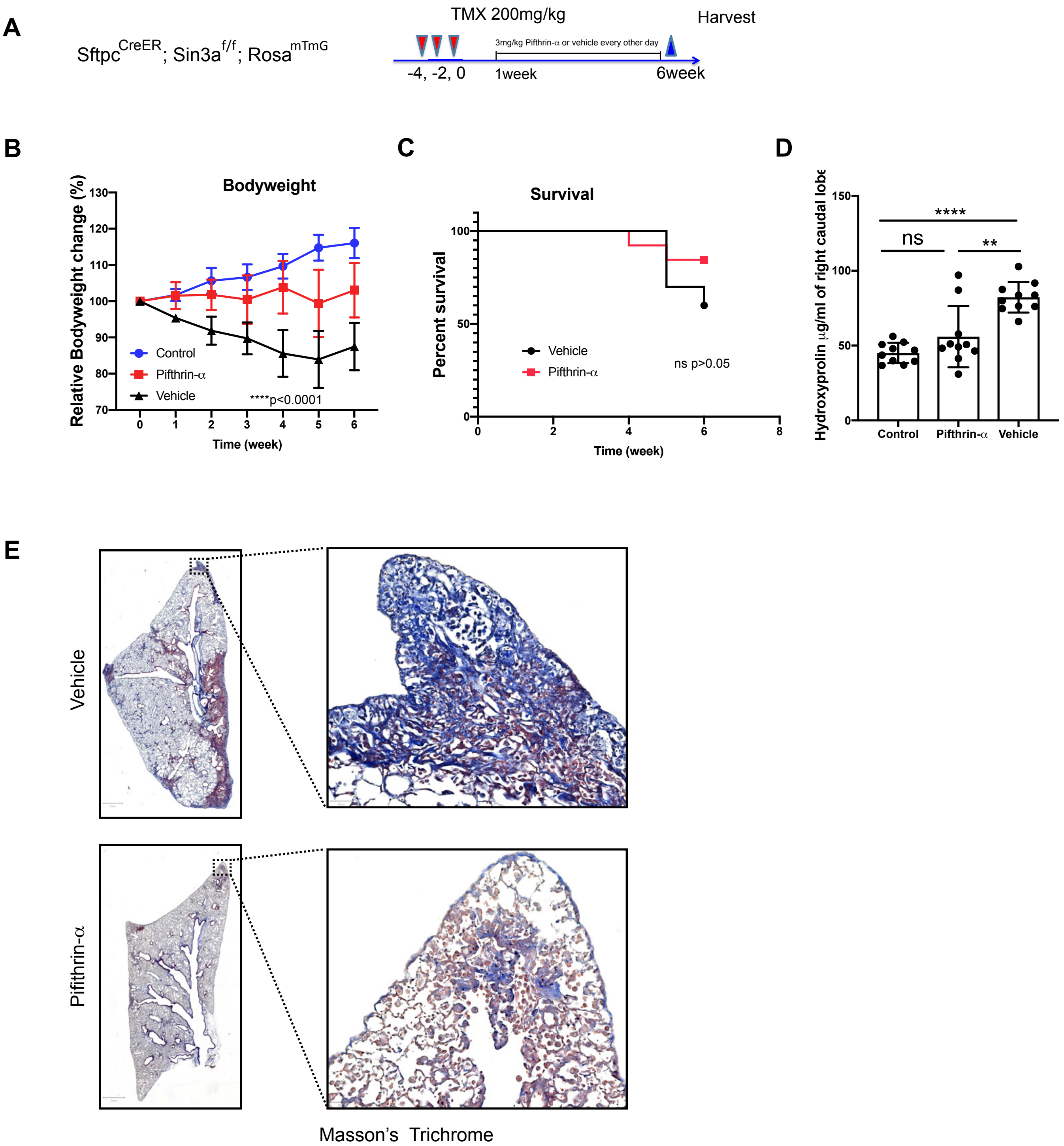


**Supplemental Figure 8. Inhibition of p53 by p53 inhibitor Pifithrin-α mitigates lung fibrosis resulting from Sin3a-LOF in AT2 cells. (Related Fig 4).**

A. Schematic outline of experiment design for p53 inhibitor Pifithrin-α treatment. B. Bodyweight change of Sin3a-LOF mice comparing Pifithrin-α treatment and vehicle treatment groups 6 weeks post-tamoxifen exposure. ****$p<0.0001$ for two-tailed student t-test comparing Pifithrin-α vs vehicle treated group at each time point since 2 weeks after tamoxifen exposure. C. Survival curve of Sin3a-LOF mice comparing Pifithrin- treatment and vehicle treatment groups 6 weeks post-tamoxifen exposure. $p>0.05$ (none significant) by Log-rank (Mantel-Cox) survival analysis. D. Hydroxyproline content in right caudal lobe of Sin3a-LOF mice treated with either Pifithrin- or vehicle, 6 weeks after tamoxifen treatment (n=10). E. Masson's trichrome staining 6 weeks post-tamoxifen treatment comparing Pifithrin- and vehicle treatment groups. p-value calculated by two-tailed student t-test. * $p<0.05$, ** $p<0.01$, *** $p<0.001$, **** $p<0.0001$.

Supplemental Figure 9.

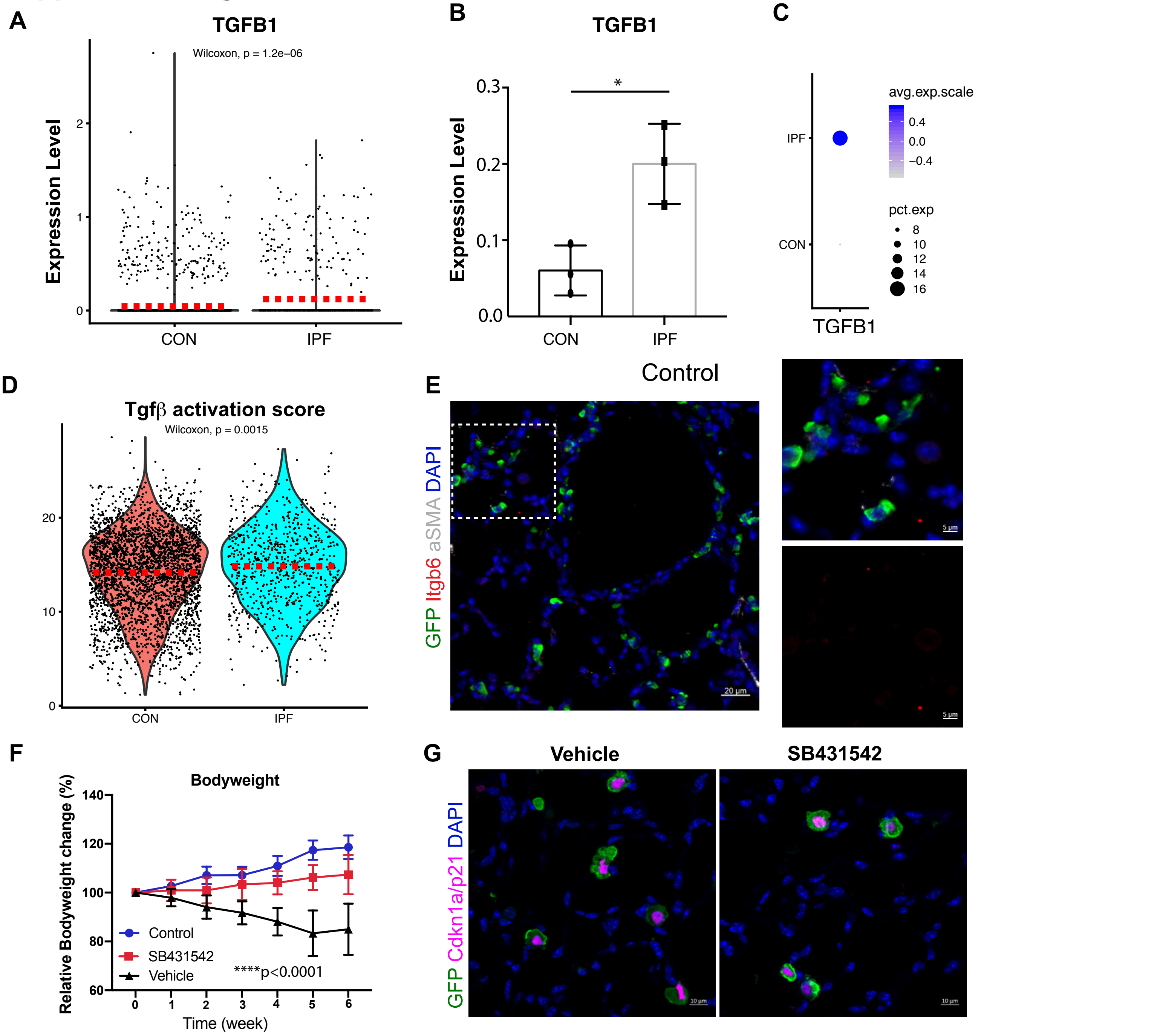


**Supplemental Figure 9. Inhibition of TGFβ mitigates lung fibrosis resulting from Sin3a-LOF in AT2 cells. (Related Fig 4 and Fig 5).**

A. Violin plot representation showing relative expression of TGFB1 in human epithelial scRNA-Seq AT2 cell subsets. B. Average expression of TGFB1 for AT2 cells of each patient sample subsets from human epithelial scRNA-Seq. C. Dotplot visualization showing relative expression of TGFB1 for AT2 cell subsets from human epithelial scRNA-Seq. D. Violin plot representation showing TGFβ activation score in human epithelial scRNA-Seq AT2 cell subsets. Dash line indicates median expression. E. Representative immunofluorescence staining of lineage reporter (GFP), Itgb6 and αSMA of control mice (SftpcCreER; RosamTmG). F. Bodyweight change of Control and SB431542 treated or vehicle treated Sin3a-LOF mice groups 6 weeks post-tamoxifen exposure. *p<0.01 and ****p<0.0001 for two-tailed student t-test comparing Pifithrin- vs vehicle treated group at 2 weeks after tamoxifen exposure and each time point 3 weeks after tamoxifen exposure respectively. G. Representative immunofluorescence staining of lineage reporter (GFP) and Cdkn1a/p21 of Vehicle control and SB431542 treated Sin3a-LOF mouse lung tissues 6 weeks post-tamoxifen treatment. Scale bar = 10μm.

#### Supplemental Figure 10.

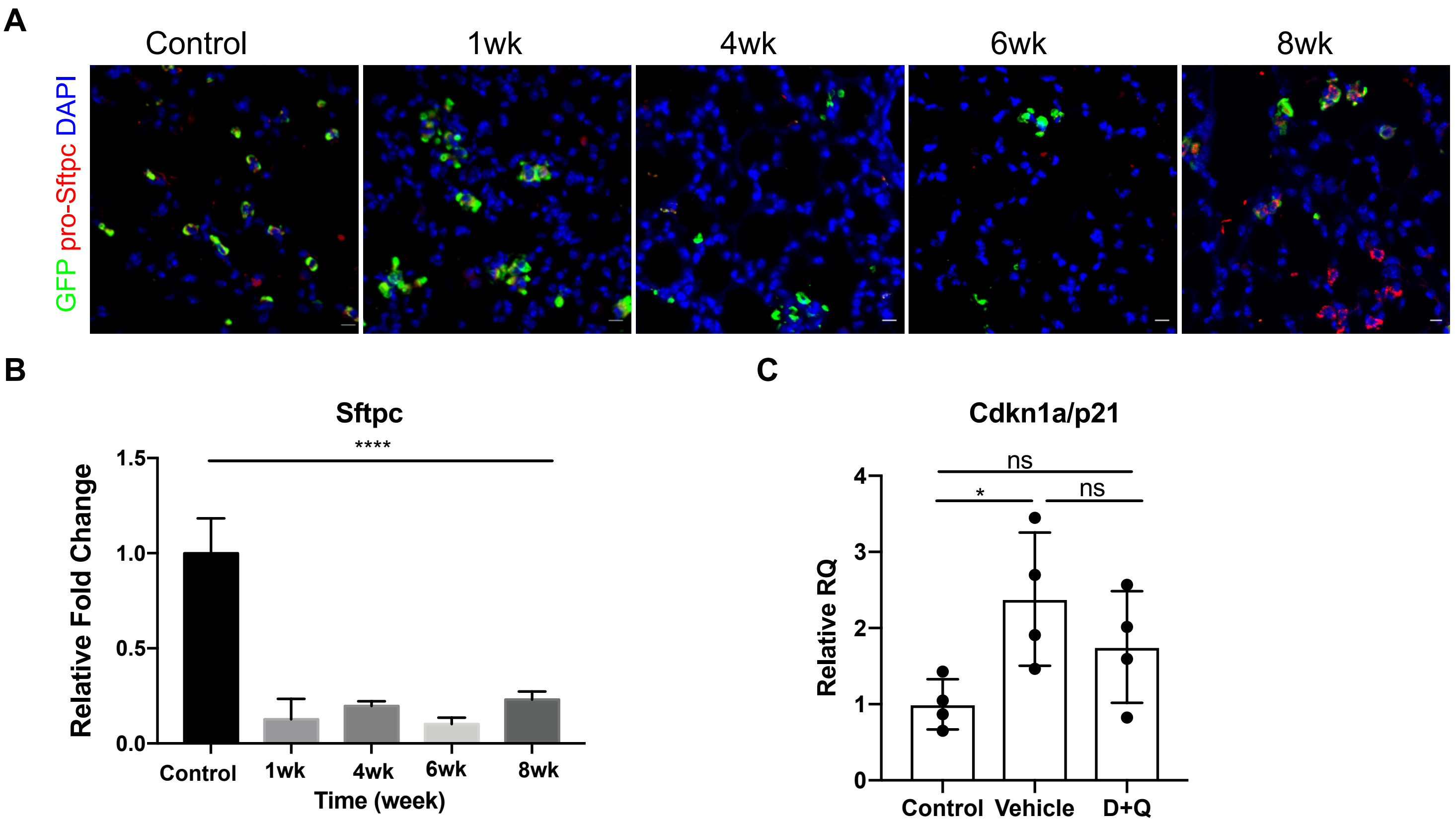


**Supplemental Figure 10. Senescence rather than loss of AT2 cells results in progressive lung fibrosis. (Related Fig 6).**

A. Representative immunofluorescence staining of lineage reporter (GFP) and Pro-Sftpc of control and DTA mouse lung tissues indicating AT2 cell ablation efficiency. Scale bar = 10μm. B. Real-time quantitative PCR of control and DTA mouse lung tissues for relative changing expression of Sftpc indicating AT2 cell ablation efficiency. C. Real-time quantitative PCR of Control, Vehicle treated and D+Q treated Sin3a-LOF mouse lung tissues for relative changing expression of Cdkn1a/p21 indicating senolytic drug treatment efficiency. p-value calculated either by two-tailed student t-test or one-way ANOVA. * $p<0.05$, ** $p<0.01$, *** $p<0.001$, **** $p<0.0001$.
